# Factors influencing patterns of gene expression in large bowel mucosa

**DOI:** 10.1101/2022.08.30.505238

**Authors:** P.G. Vaughan-Shaw, L.Y. Ooi, M. Timofeeva, G. Grimes, F.V.N. Din, S.M. Farrington, M.G. Dunlop

## Abstract

**Aim:** Variation in genomic sequences that control gene expression in target tissues is increasingly recognized as the underlying basis of disease susceptibility, especially cancer. This is particularly relevant to common genetic variation impacting on cancer risk. Exogenous factors may also influence gene expression, potentially leaving a transcriptomic signature of environmental risk factors in normal tissues. Therefore, understanding endogenous and exogenous influences over gene expression patterns in normal tissue is critical to the study of functional genomics and its relationship to disease susceptibility. Here, we investigated demographic and sampling variables that could impact on gene expression in normal colorectal mucosa.

**Method:** We prospectively collected normal mucosa from 424 patients undergoing colorectal surgery or outpatient assessment through surgical stripping of normal mucosa from resected colorectal specimens or rectal biopsy. Gene expression was assessed using Illumina HT-12 microarrays and analysed against demographic (age, gender, BMI) and sampling factors (general anaesthesia, cleansing bowel agents, sample site, time to RNA preservation) using adjusted linear regression modelling.

**Results:** Age, gender, smoking status, sampling under anaesthetic, sample site and sampling method were associated with differential gene expression in the adjusted model. BMI or use of cleansing bowel preparation did not impact gene expression. Age was associated with differential expression of 16 genes and significant enrichment in pathways relevant to tumourigenesis, including immune process, cell proliferation, adhesion and death. Sample site was associated with differential expression of 2515 genes, with 1102 genes more highly expressed in the proximal colon (proximal to splenic flexure). Gender and sampling under anaesthetic were associated with differential expression of 99 and 851 genes respectively. Increased time to RNA preservation (45-90 minutes) was associated with enrichment of pathways consistent with tissue ischaemia including *‘response to wounding’, ‘apoptotic process’*, and *‘response to oxygen levels’*.

**Conclusions:** Demographic and sampling factors influence gene expression patterns in normal colorectal mucosa, often with a large magnitude of effect. Meanwhile, greater time to RNA preservation is associated with patterns of gene expression consistent with tissue ischaemia which questions the generalisability of assessment of gene expression patterns generated from post-mortem studies. These results highlight the importance of fully adjusted expression analyses and may indicate mechanisms underlying age, gender and site-specific differences in CRC incidence, progression and outcome.

## Introduction

Methods of whole transcriptome profiling, including DNA microarray and RNA sequencing are increasingly available and frequently used in studies of *in vivo* gene expression. Understanding the influence of baseline demographic factors on expression is required for appropriate adjustment of these data to reduce bias but may also define mechanisms underlying variation in disease incidence or outcome. Significant variation in sampling and processing methods exist in human expression studies published to date with cadaveric tissue, operative samples, endoscopic biopsies, formalin-fixed paraffin-embedded tissue and isolated cells all being used [1-9] yet the differential impact of these methods expression is not known.

Colorectal cancer is common, with over 40,000 new cases in the UK each year. High disease incidence, the availability of redundant tissue from resected colonic specimens and the relative ease of endoscopic sampling has led to numerous gene expression studies in colorectal cancer with differential expression commonly reported in CRC and adenoma tissue [10]. Such data could provide valuable insight into the aetiology of colorectal cancer and inform preventive and therapeutic strategies while improving personalised approaches to the disease. However, individual studies tend to be small, and so important findings may only be uncovered by assimilating data from different studies. This approach requires a firm understanding of the impact of common clinical, demographic and sampling factors on gene expression. Such analysis might also provide novel insight into the mechanism of colorectal tumourigenesis, given marked gender and age-specific variation in incidence. Meanwhile, site-specific variation in CRC incidence [11], biology [12, 13], response to adjuvant therapy [14], and prognosis [15-18] are well recognised, yet incompletely understood. Differences may reflect different embryological origin, different exposure to faecal stream, microbiome and carcinogens [19] or differential genetic influences over colorectal cancer risk and gene expression [20].

Colorectal tissue is most commonly sampled during colorectal resection surgery or colonoscopy. However, both procedures require a period of pre-procedure starvation which may influence gut transit, systemic glucose levels, gut microbiota and consequently mucosal gene expression. Meanwhile, colonoscopy and certain surgical procedures require bowel preparation with strong laxatives, while resection surgery requires general anaesthesia-factors which likely impact mucosal gene expression yet have not been systematically investigated to date. Expression in cadaveric tissue may overcome such issues [9] yet is likely prone to the effects of tissue ischaemia, which remain largely unexplored in the context of colorectal gene expression.

In this study we systematically investigated the impact of common demographic and sampling factors on gene expression in normal colorectal mucosa.

## Method

### Study population

Participants were sampled from two studies. The Study of Colorectal Cancer in Scotland (SOCCS) is a prospective, population-based case-control study designed to identify genetic and environmental factors that have an impact on CRC risk and survival outcome [21]. All SOCCS participants with available normal mucosa for gene expression analysis were included. The Scottish Vitamin D study (SCOVIDS) was designed and initiated to enable the recruitment and sampling of subjects not undergoing major colorectal resection. All participants provided informed written consent, and research was approved by local research ethics committees (13/SS/0248; 11/SS/0109 and 01/0/05) and National Health Service management (2014/0058; 2013/0014 and 2003/W/GEN/05). Clinical and sampling variables were collected from patient case notes and pathology records, entered into a prospective study database and extracted for analysis.

### Mucosa sampling and storage

Normal colorectal mucosa (NM) was sampled from resected surgical specimens or with rectal biopsy. For resection samples, mucosa was surgically stripped from the submucosa and immersed in the stabilization solution RNAlater (Invitrogen). This process was commonly performed by a local tissue bank with intervals of 45-90 minutes between surgical resection and RNAlater immersion. A subset of stripped mucosa samples were taken by research fellow (PVS), with RNA preservation in RNAlater within 5 minutes of surgical resection. Subjects sampled in the clinic or during minor surgical procedures underwent rigid sigmoidoscopic rectal biopsy, with samples immediately placed in RNAlater. All samples were kept in RNAlater for 24-72 hours prior to RNA extraction or storage at -80□C. To facilitate an assessment of biological variation, a separate NM sample was taken from the same site, at the same time from 24 subjects.

### Assessment of gene expression

NM samples were chopped into ∼3mm^3^ pieces, immersed in 1ml TRIzol (Applied Biosystems) and homogenised using a standard mechanical tissue lyser at 50Hz using 7mm RNase free ball bearings. RNA was extracted and purified from NM using a proprietary RNA extraction kit (Ribopure kit, Applied Biosystems) according to the manufacturer’s protocol. RNA was quantified by NanoDrop (NanoDrop Technologies) and yield and integrity was assessed using the Agilent 2100 Bioanalyzer.

Gene expression was primarily assessed using DNA microarray. Total RNA was converted to double-stranded cDNA, followed by in vitro transcription amplification to generate labelled cRNA (Illumina TotalPrep™ RNA Amplification Kit). Gene expression profiling was undertaken using the HumanHT-12 v4.0 Expression BeadChip Arrays (Illumina) and IScan NO660 scanner, providing coverage of 47,231 transcripts and >31,000 annotated genes derived from the National Centre for Biotechnology Information Reference Sequence RefSeq Release 38 (November 7, 2009). To assess technical variation analysis of 18 RNA samples was repeated and expression data compared.

Microarray data was exported from BeadStudio (Illumina) and processed in R using the *limma* package. In brief, the steps involved were background correction using negative controls, quantile normalization to remove technical variation and finally log2 transformation. Control probes and probes that were not expressed in at least three arrays to a detection-value of ≥5% were excluded. A standard approach to batch correction using ComBat was performed to control for batch effects [22].

### Statistical analysis

All statistical analysis was undertaken in R [23]. Expression values for biological and technical replicate samples were averaged at outset except for assessment of respective variation in expression. Investigation of differential gene expression was undertaken using the *lmFit* and *eBayes* functions within the limma package. Factors tested included age, gender, BMI, smoking status, sampling under general anaesthetic, sampling following bowel cleansing (i.e. oral laxative or phosphate enema administered per rectum), sample side (i.e. proximal or distal to the splenic flexure), and sample method (biopsy or stripped mucosa). Adjustment for multiple testing was undertaken and FDR p-values derived [24]. Linear regression modelling was used to adjust for relevant clinical and sampling variables.

Assessment of technical and biological variation in gene expression was performed by comparison of inter-sample Pearson correlation coefficient using global HT12 expression data.

### Pathway analysis

Gene ontology and enrichment analysis was undertaken using the ‘GOrilla’, Gene Ontology enRIchment anaLysis and visuaLizAtion tool [25]. Process ontologies were investigated using gene lists ranked by p value.

## Results

466 NM samples from 424 subjects were included in analysis (Table 1, Supplementary Table 1).

**Table 1.** Baseline characteristics, sampling variables and vitamin D status in participants included in observational analysis. Other diagnoses included anal skin tag (n=2), idiopathic constipation, Familial adenomatous polyposis coli, Hidradenitis suppurativa and Peutz-jeghers (all n=1). Sample method applies to unique patient, ignoring technical and biological replicate samples.

|  |  |
| --- | --- |
| **Age** | Median 67 (range 17-91) |
| **Gender** | 202 (48% female) |
| **Diagnosis** |  |
| Colorectal cancer | 274 |
| None (healthy) | 59 |
| Haemorrhoids | 20 |
| Past medical history of CRC | 13 |
| Diverticular disease | 12 |
| Colorectal adenoma | 10 |
| Fistula-in-ano | 7 |
| Anal intra-epithelial neoplasia | 5 |
| Pelvic tumour (not colorectal) | 5 |
| Fissure-in-ano | 4 |
| Small bowel tumour | 3 |
| Anorectal fibroepithelial polyp | 3 |
| Pilonidal disease | 3 |
| Other | 6 |
| **Sample site** |  |
| Caecum | 14 |
| Transverse colon | 95 |
| Not specified (proximal) | 3 |
| Not specified (distal) | 17 |
| Descending colon | 117 |
| Sigmoid colon | 17 |
| Rectum | 161 |
| **Bowel preparation** | 155 |
| Phosphate enema | 99 |
| Picolax (Sodium Picosulfate) | 38 |
| Moviprep (polyethylene glycol electrolyte sln) | 16 |
| Yes, not specified | 2 |
| **Sampled under general anaesthetic** | 342 (81%) |
| **Sample method** |  |
| Stripped mucosa | 298 |
| Biopsy | 126 |

Technical replicate samples (i.e. expression analysis of the same RNA sample, n=18) and biological replicate samples (i.e. RNA extraction from different piece of NM sample, n=24) showed high intra-patient correlation of gene expression (technical R=0.98, biological R=0.97). Age, gender, smoking status, sampling under anaesthetic, sample site (proximal or distal colorectum), and sampling method were associated with differential gene expression in the adjusted model (Table 2).

**Table 2.** Clinical and sampling factors and association with probe expression in adjusted linear regression model. Adjusted linear regression model performed adjusting for cancer status, sampling method, sample side, anaesthetic, age and/ or gender. Est – linear regression estimate,. BMI body mass index. Significant up-or down-regulation was defined as FDR <0.05. Bowel preparation refers to any pre-operative bowel cleansing agent used. Distal colorectum taken as any site distal to anatomical splenic flexure. Collinearity existed between sampling site, sampling method and CRC status, with cancer-free subjects sampled predominantly by biopsy of rectum (i.e. distal colorectum), and samples from proximal colorectum all stripped mucosa samples, predominantly from CRC patients (see supplementary file).

| Factor | Category comparison | Up-regulated | Down-regulated |
| --- | --- | --- | --- |
| Age | Increasing years | 5 | 11 |
| Gender | Male vs. Female | 45 | 54 |
| BMI | Increasing BMI | 0 | 0 |
| Anaesthetic | Yes vs. No | 601 | 250 |
| Bowel preparation | Yes vs. No | 0 | 0 |
| Sample side | Proximal vs. distal | 1102 | 1413 |
| Sample method | Stripped mucosa vs. biopsy | 8 | 6 |
| Processing time | RNA later >60min vs.<10min | 0 | 1 |

### Age

Age was significantly associated with the expression of 16 genes, with increased age associated with higher expression in 5 genes (Table 3, Supplementary Figure 1), and 38 significantly enriched processes on pathway analysis (Supplementary Table 2). Several processes relevant to tumourigenesis were enriched, including terms relating to immune processes, cell proliferation, adhesion and death. Ten probes were differentially expressed when age was considered as a binary factor (<>67 years), validating 6 of the hits with age as a continuous variable (Supplementary Table 3).

**Table 3.** Differentially expressed probes by increasing subject age. Adjusted linear regression model performed adjusting for cancer status, sampling method, sample side, anaesthetic and gender. Est – linear regression estimate.

| Probe ID | Gene | Est. | FDR |
| --- | --- | --- | --- |
| ILMN_1655765 | MRPS21 | -0.012 | 1.27e-11 |
| ILMN_1841495 | LHFPL4 | 0.005 | 2.69e-08 |
| ILMN_2224486 | C3orf14 | -0.012 | 2.69e-08 |
| ILMN_1699665 | CLIC6 | -0.026 | 4.58e-08 |
| ILMN_1660292 | MRPS21 | -0.012 | 6.49e-08 |
| ILMN_1812824 | SST | -0.031 | 4.39e-06 |
| ILMN_1751445 | PPY | 0.018 | 8.21e-06 |
| ILMN_1741597 | LOC650200 | -0.009 | 1.84e-05 |
| ILMN_1783247 | C10orf1 | 0.010 | 9.17e-05 |
| ILMN_1682449 | ZNF518B | -0.008 | 0.0002 |
| ILMN_1691290 | CELSR3 | 0.0085 | 0.000206 |
| ILMN_1777318 | C9orf64 | -0.007 | 0.003567 |
| ILMN_1758173 | TMEM99 | -0.0067 | 0.004595 |
| ILMN_1685608 | NPTX2 | -0.0104 | 0.006612 |
| ILMN_2367487 | ABHD12B | -0.0078 | 0.007225 |
| ILMN_1796417 | ASNS | 0.0067 | 0.049035 |
| ILMN_1745501 | DNALI1 | -0.0081 | 0.049035 |

### Gender

Gender was associated with differential expression of 99 genes, with higher expression of 45 genes in Males, and lower expression of 54 genes (Supplementary Table 4). Top ranked hits were commonly Y-linked gene (e.g. *EIF1AY, RSP4Y1*) more highly expressed in males, while the X-inactive specific transcript (*XIST*) gene responsible for silencing of the inactive X chromosome showed significantly greater expression in females (Table 4). Seventeen enriched GO terms were identified associated with gender, including terms relevant to tumourigenesis such as ‘oxidation-reduction process’, *‘immune response’* and *‘methylation’* (Supplementary Table 5).

**Table 4.** Top ten ranked differentially expressed probes by gender in adjusted analysis. Adjusted linear regression model performed adjusting for cancer status, sampling method, sample side, and age. FC-estimated fold-change calculated from linear regression coefficient.

| Probe ID | Gene | FC | FDR |
| --- | --- | --- | --- |
| ILMN_3238417 | LOC100133662 | 38.05 | 3.91E-279 |
| ILMN_1755537 | EIF1AY | 22.32 | 6.76E-277 |
| ILMN_1783142 | RPS4Y1 | 79.34 | 1.92E-275 |
| ILMN_1670821 | CYorf15A | 8.69 | 2.81E-266 |
| ILMN_2228976 | EIF1AY | 4.99 | 4.15E-248 |
| ILMN_1685690 | JARID1D | 8.06 | 4.38E-243 |
| ILMN_1764573 | XIST | -10.06 | 9.14E-232 |
| ILMN_2191331 | RPS4Y2 | 7.21 | 3.08E-210 |
| ILMN_1756506 | CYorf15B | 3.97 | 1.30E-169 |
| ILMN_3212921 | LOC643123 | 3.34 | 7.30E-167 |

**Table 5.** Top 10 ranked differentially expressed probes in samples taken from proximal versus distal colorectum in adjusted analysis. Adjusted linear regression model performed adjusting for cancer status, sampling method, age and gender. FC-estimated fold-change calculated from linear regression coefficient.

| Probe ID | Gene | FC | FDR |
| --- | --- | --- | --- |
| ILMN_1801832 | *PRAC* | -30.70 | 4.46e-79 |
| ILMN_3248384 | *PRAC* | -29.86 | 4.56e-59 |
| ILMN_2391400 | *PITX2* | 4.72 | 3.37e-44 |
| ILMN_1796847 | *PITX2* | 2.33 | 8.79e-30 |
| ILMN_1742677 | *HOXB13* | -4.56 | 1.77e-29 |
| ILMN_2204545 | *ST3GAL4* | -2.71 | 1.77e-29 |
| ILMN_1691736 | *ST6GALN*<br>*AC6* | -2.41 | 2.84e-27 |
| ILMN_1769839 | *L1TD1* | 2.58 | 3.21e-27 |
| ILMN_1746676 | *CLDN8* | -3.86 | 2.56e-26 |
| ILMN_3248309 | *LOC7322*<br>*15\** | -1.54 | 5.28e-26 |

### General anaesthesia and bowel cleansing agents

Sampling of normal mucosa while the subject was under general anaesthesia was associated with differential expression of 851 genes (Top hit *IRX2*, FC 1.85, p=3.14e-10, Supplementary Table 6), and enrichment of 410 GO terms (Supplementary Table 7). No probe was significantly associated with pre-operative bowel preparation (top hit *NPTX2*, FDR p=0.58), although significant collinearity was present, with all subjects sampled after bowel preparation also sampled under general anaesthesia.

**Table 6.** Probes with expression associated with sampling method in adjusted model. Adjusted linear regression model performed adjusting for cancer status, sample side, age and gender. FC-estimated fold-change calculated from linear regression coefficient.

| Probe ID | Gene | FC | Adjusted p-value |
| --- | --- | --- | --- |
| ILMN_1776125 | *TRK1* | -1.36 | 0.001 |
| ILMN_1711078 | *CDC2L2* | 1.48 | 0.008 |
| ILMN_1742332 | *KCTD12* | 1.45 | 0.023 |
| ILMN_2391400 | *PITX2* | -1.64 | 0.024 |
| ILMN_2330552 | *CDC2L2* | 1.41 | 0.024 |
| ILMN_1801832 | *PRAC* | 2.40 | 0.025 |
| ILMN_3248384 | *PRAC* | 2.37 | 0.027 |
| ILMN_1666109 | *MB* | -1.47 | 0.030 |
| ILMN_1708414 | *GNL3L* | 1.54 | 0.042 |
| ILMN_1696870 | *TGFBRAP1* | 1.28 | 0.042 |
| ILMN_1743506 | *CCDC137* | 1.10 | 0.042 |
| ILMN_1763436 | *SETX* | 1.28 | 0.043 |
| ILMN_1873075 | *Unknown* | 1.12 | 0.043 |
| ILMN_1786920 | *JARID1A* | 1.20 | 0.045 |
| ILMN_2383455 | *SUOX* | -1.23 | 0.045 |
| ILMN_1726547 | *MAP3K5* | 1.24 | 0.046 |

### Sample side

Sample side was associated with differential expression of 2515 genes, with 1102 genes more highly expressed in the proximal colon (proximal to splenic flexure), and 1413 genes upregulated in the distal colorectum as previously reported [20]. The top hit (*PRAC*), showed ∼30-fold greater expression in distal colorectum, with 47 genes in total demonstrating at least 2-fold difference in expression (Supplementary Table 8). qRT-PCR validation was undertaken for a subset of samples (39 proximal and 76 distal colon samples, see [20]), for 3 of the top ranked genes confirming significant differential expression in *PRAC* (*p*<2.2e-16), *PITX2* (*p*<2.2e-16) *and L1TD1* (*p*=5.3e-13). Highly significant correlations with Spearman Rho values of >0.75 were observed between the two different expression quantification techniques [20].

### Sample method

In the current study 126 samples were taken by biopsy of the rectal mucosa, while 298 samples were stripped mucosa taken from resected colorectal specimens, with 14 differentially expressed genes dependent on sample method (Table 6). To assess for difference in amount of stromal tissue within the respective mucosal samples the expression of 10 genes with known stromal/ epithelial signatures was assessed. There was no evidence of differential expression in these genes known to be more highly expressed in stromal tissue (*CD93, PCDH18, VIM, ABCA8, GJA1, ACTA2, POSTN)* or epithelial tissue (*EPCAM, CCND1*, Supplementary Table 9). Finally, matched biopsy and stripped mucosa sampling of the same site from 9 resected colorectal specimens revealed no difference in expression of 4 candidate genes (*EPCAM, KRT18, DES, VIM*, qRT-PCR analysis, Supplementary Figure 5).

### Time to RNA preservation

Immediate RNA preservation by immersion in RNAlater was performed for 165 samples, while in the remainder RNA preservation in RNAlater or by snap freezing occurred between 45-90 minutes following resection/ biopsy. A single gene was significantly downregulated in samples where RNA preservation occurred beyond 45 minutes (*TSC22D3*, FC -1.66, FDR p=0.006). Gene ontology analysis revealed 72 significantly enriched terms associated with interval RNA preservation, many consistent with the presence of anaerobic metabolism, ischaemia and cell death (Supplementary Table 10, Figure 1)

**Figure 1.**
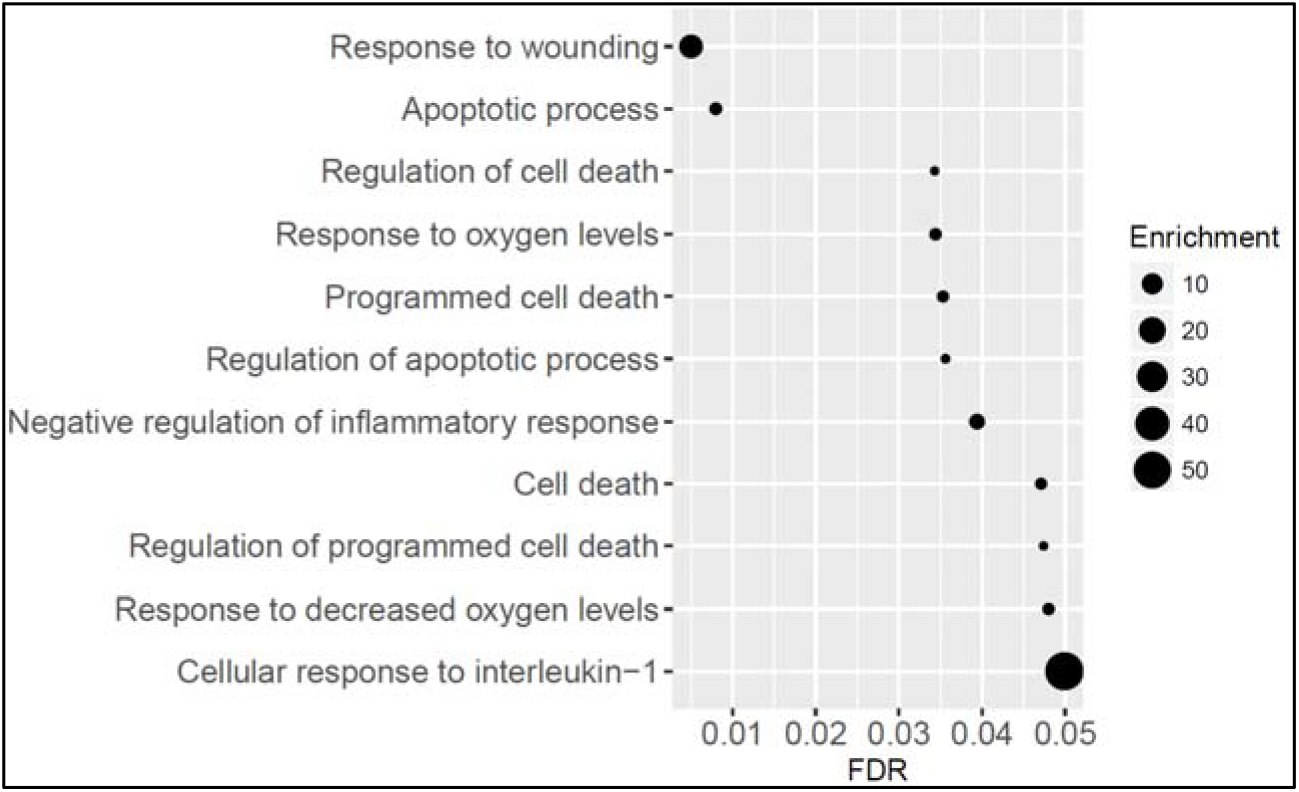
GO enrichment of selected terms in samples with time to RNA preservation >45 minutes. Differential expression was calculated using adjusted linear regression model and genes ranked by p value entered into the GO enrichment software GOrilla. FDR q-value’ is the correction of the above p-value for multiple testing using the Benjamini and Hochberg method [9]. For the ith term (ranked according to p-value) the FDR q-value is (p-value * number of GO terms) / i. Enrichment = (the number of genes in the intersection/ number of genes in the top of the input list)/(total number of genes associated with a specific GO term/ total number of genes).

## Discussion

In this prospective study of normal mucosa samples from 424 human subjects we show that demographic and sampling factors significantly impact gene expression in the human colorectum, often with a large magnitude of effect. Differential expression relating to age, gender and sample site highlight the need to adjust for these factors in expression analyses, while such differences may inform the understanding of variation in CRC incidence, progression and outcome dependent on such factors. Evidence that a modest delay to RNA preservation following surgical resection results in pathway enrichment consistent with tissue ischaemia questions the use of post-mortem tissue samples in expression studies.

Comparative expression studies of colorectal tissue offer the promise of improved understanding of the carcinogenesis process, discovery of biomarkers for screening and profiling and the identification of novel preventive strategies and therapeutic targets. Numerous expression studies relating to CRC have been published in the last two decades [10, 26], yet individual studies are invariably small, with few including more than 50 patients. Synthesis of data from individual studies may unmask important results, as is demonstrated in GWAS meta-analyses. However, unlike GWAS where genetic variants are constant, expression is a variable phenotype dependent not only on cancer status, but also demographic factors including diet, age and gender, and further influenced by sampling factors. To date this has limited attempts to synthesise colorectal expression data to simple overlap analysis [10]. To facilitate future synthesis of individual studies, the current study provides data to indicate which baseline variables should be recorded in studies of gene expression and suggests that failure to adjust for such factors would result in significant bias.

Identification of age, gender and site-specific gene expression may add to the current understanding of the aetiology of CRC given known differences in incidence, molecular subtype and outcomes dependent on such factors. As with other common cancers, CRC is primarily an age-related disease. Previous studies demonstrate changes in colonic mucosa DNA methylation with aging [27], while aberrant methylation of promoter regions is associated with inactivation of tumor-suppressor genes in neoplasia [28]. Indeed, altered DNA methylation is seen in a considerable proportion of age-related gene promoters in CRC or adenoma tissue [29] which likely contributes to increased CRC incidence with age. Two such genes which are reported to be hypermethylated in both adenoma and CRC tissue when compared to normal mucosa demonstrated lower expression with increased age in the current study-*SFRP1* a *Wnt* antagonist coding Secreted frizzled related protein 1 (p=0.0005)[30, 31] and *DKK3*, coding Dickkopf *WNT* signaling pathway inhibitor 3 (p=0.02), supporting the concept that changes in gene expression with age drives tumourigenesis. Meanwhile, the large number of differentially expressed genes between the proximal and distal colorectum and enriched processes on pathway analysis may provide valuable insight into previously reported site-specific differences in CRC incidence [11], biology [12, 13], response to adjuvant therapy [14], and prognosis [15-18]. The cause of such differences is not fully understood but may reflect different embryological origin or different exposure to faeces and carcinogens [19]. We have previously described topographical differences in genomic control over gene expression relevant to CRC risk which may underlie site-specific variation in CRC and could, in time, inform individualised CRC screening programmes or site-specific adjuvant/ neo-adjuvant CRC therapy.

The final part of this study looks at sampling factors including method of sampling and processing, with evidence of tissue ischaemia after a modest delay to RNA preservation. This data questions the validity of gene expression analysis of cadaveric tissue performed in expression studies including the GTEX study [9, 32]. In the current project expression in tissue samples immediate immersed in RNAlater was compared to samples where RNA preservation occurred up to 90 minutes following extirpation of the surgical specimen. Even this short delay led to significant enrichment of processes linked to tissue ischaemia, yet in the GTEX cadaveric tissue expression study the interval from death to sample collection for colonic tissue was 236.5 minutes. In GTEX, prolonged post-mortem interval (PMI) is associated with reduced RNA integrity number value (RIN) [33], yet impact on expression is not reported, while in other published studies significant changes in colonic gene expression are seen after 40-60 minutes at room temperature [34, 35]. These data suggest that results from cadaveric tissue should be treated with caution and where possible tissue collection procedures should ensure snap freezing or RNA preservation within one hour.

The current study is a pragmatic study of gene expression in normal colorectal mucosa collected by clinical research fellows and the local tissue bank. While the heterogeneity in sampling factors may be seen as a limitation, it has allowed an assessment of the impact of such factors on expression. Meanwhile, inherent collinearity exists between certain factors which has precluded investigation of such factors and expression. For example, the vast majority of NM samples from cancer-free subjects were taken from the rectum and so a rigorous investigation of cancer status on gene expression is not possible. To address this, future efforts should include pre-operative rectal biopsy sampling from a cohort of patients with CRC cohort. Meanwhile, the absence of significant associations between bowel preparation and gene expression in the current dataset, supports utilisation of colonoscopic biopsies as a means of including large numbers of proximal colonic samples from both case and control subjects.

In conclusion, the current study is the largest descriptive expression analysis of normal colorectal mucosa to date. Marked gender, age, site and sampling-specific expression differences are seen, indicating both the importance of fully adjusted analyses and also the potential value of differential expression data in the investigation of colorectal tumourigenesis. Meanwhile, the impact of sampling factors, in particular time to RNA preservation, on expression highlights the importance of streamlined tissue collection and processing procedures.

## Data Availability Statement

Expression data available at https://pubmed.ncbi.nlm.nih.gov/33937989/

## Acknowledgements

We acknowledge the excellent technical support from Marion Walker and Stuart Reid. We are grateful to Donna Markie and Fiona McIntosh, and all those who continue to contribute to recruitment, data collection, and data curation for the Study of Colorectal Cancer in Scotland studies. We acknowledge that these studies would not be possible without the patients and surgeons who take part and the NHS Lothian Bioresource team which contributed to the collection and storage of NM samples for this study. We acknowledge the expert support on sample preparation from the Genetics Core of the Edinburgh Wellcome Trust Clinical Research Facility. This research has been conducted using the UK Biobank Resource under Application Number 7441.

