## Supplementary Tables and Figures for "Factors influencing patterns of gene expression in large bowel mucosa"

Supplementary Table 1 Tumour distribution in included participants with CRC

| CRC site | PMH CRC | Current CRC |
| --- | --- | --- |
| Right colon not otherwise specified | 0 | 1 |
| Caecum | 2 | 55 |
| Ascending colon | 4 | 32 |
| Hepatic flexure | 0 | 7 |
| Transverse colon | 0 | 20 |
| Splenic flexure | 1 | 11 |
| Descending colon | 0 | 10 |
| Sigmoid colon | 6 | 63 |
| Rectosigmoid | 0 | 15 |
| Rectum | 1 | 61 |

Two patients had synchronous tumours, i.e. CRC at 2 sites, hence the total number of sites is 289 from 287 patients.

Supplementary Figure 1 Plot of top ranked probe against age

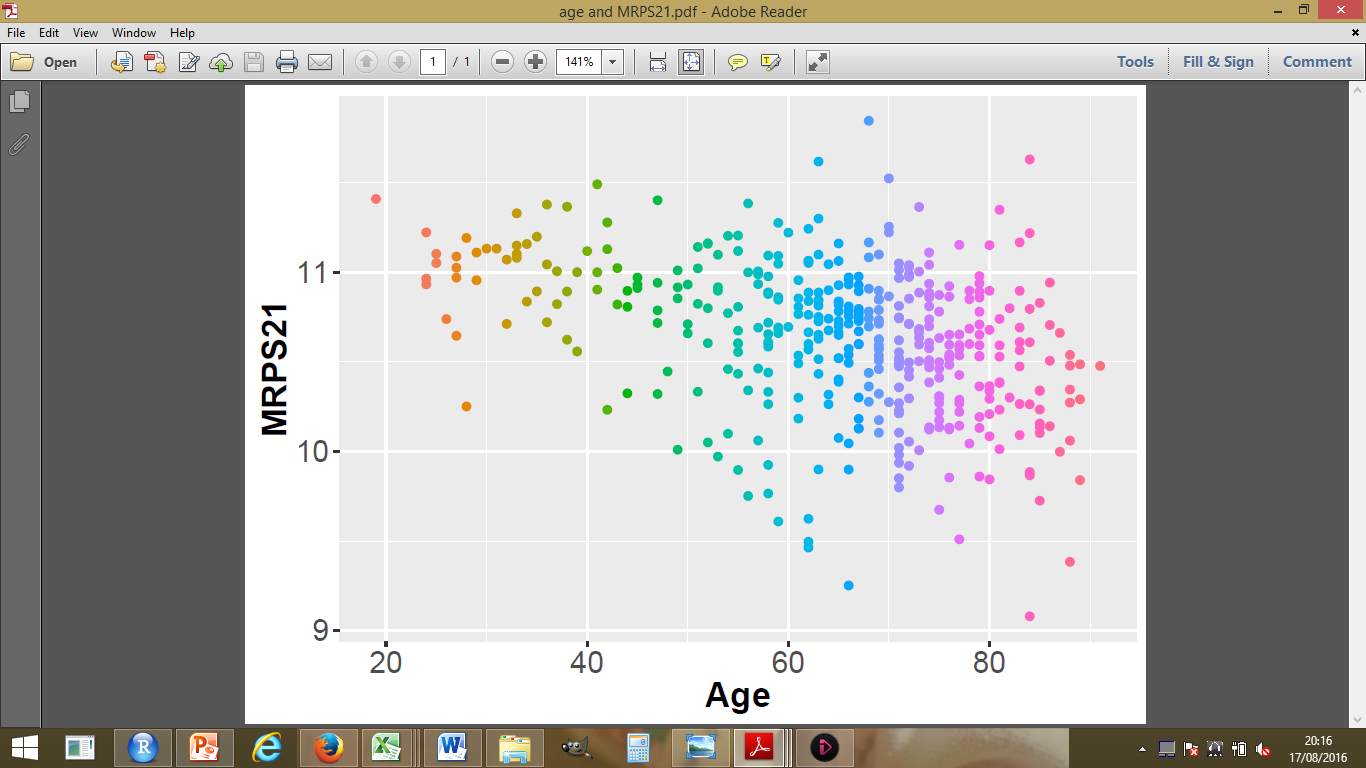

Supplementary Table 2 Enriched GO terms associated with higher age

| GO term | **Description** | **p-value** | **FDR q** |  | **Enrichment** | **Genes** |
| --- | --- | --- | --- | --- | --- | --- |
| GO:0030198 | Extracellular matrix organization | 9.20e-13 | 1.36e-08 |  | 2.55 | 70 |
| GO:0043062 | Extracellular structure organization | 9.20e-13 | 6.80e-09 |  | 2.55 | 70 |
| GO:0002376 | Immune system process | 7.30e-08 | 3.60e-04 |  | 1.47 | 204 |
| GO:0070268 | Cornification | 4.45e-07 | 1.65e-03 |  | 2.76 | 33 |
| GO:0008285 | Negative regulation of cell proliferation | 4.70e-07 | 1.39e-03 |  | 1.64 | 115 |
| GO:0051607 | Defense response to virus | 6.43e-07 | 1.58e-03 |  | 2.96 | 29 |
| GO:0012501 | Programmed cell death | 8.21e-07 | 1.73e-03 |  | 1.56 | 134 |
| GO:0009615 | Response to virus | 8.68e-07 | 1.61e-03 |  | 2.51 | 38 |
| GO:0008219 | Cell death | 9.07e-07 | 1.49e-03 |  | 1.54 | 139 |
| GO:0030199 | Collagen fibril organization | 1.28e-06 | 1.89e-03 |  | 4.42 | 16 |
| GO:0060337 | Type I interferon signaling pathway | 1.39e-06 | 1.87e-03 |  | 4.48 | 17 |
| GO:0022610 | Biological adhesion | 1.87e-06 | 2.31e-03 |  | 1.69 | 98 |
| GO:0007155 | Cell adhesion | 2.55e-06 | 2.91e-03 |  | 1.69 | 97 |
| GO:0032963 | Collagen metabolic process | 4.45e-06 | 4.70e-03 |  | 3.17 | 23 |
| GO:0050793 | Regulation of developmental process | 8.46e-06 | 8.34e-03 |  | 1.35 | 233 |
| GO:0042127 | Regulation of cell proliferation | 1.22e-05 | 1.13e-02 |  | 1.36 | 213 |
| GO:0044259 | Multicellular organismal macromolecule metabolic process | 1.52e-05 | 1.32e-02 |  | 2.97 | 23 |
| GO:0030574 | Collagen catabolic process | 1.52e-05 | 1.25e-02 |  | 3.24 | 20 |
| GO:0019221 | Cytokine-mediated signaling pathway | 1.93e-05 | 1.50e-02 |  | 1.81 | 64 |
| GO:0050678 | Regulation of epithelial cell proliferation | 2.14e-05 | 1.58e-02 |  | 2.51 | 30 |
| GO:0006955 | Immune response | 2.22e-05 | 1.57e-02 |  | 1.61 | 96 |
| GO:0008544 | Epidermis development | 2.62e-05 | 1.76e-02 |  | 2.64 | 26 |
| GO:0007044 | Cell-substrate junction assembly | 2.72e-05 | 1.75e-02 |  | 3.53 | 16 |
| GO:0035759 | Mesangial cell-matrix adhesion | 3.31e-05 | 2.04e-02 |  | 172. | 92 |
| GO:0002474 | Antigen processing and presentation of peptide antigen via MHC class I | 3.54e-05 | 2.09e-02 |  | 3.90 | 16 |
| GO:0048856 | Anatomical structure development | 3.76e-05 | 2.14e-02 |  | 1.56 | 99 |
| GO:0002252 | Immune effector process | 4.01e-05 | 2.20e-02 |  | 1.62 | 89 |
| GO:0009888 | Tissue development | 4.60e-05 | 2.43e-02 |  | 1.58 | 91 |
| GO:0048869 | Cellular developmental process | 4.85e-05 | 2.48e-02 |  | 1.63 | 83 |
| GO:2000026 | Regulation of multicellular organismal development | 5.28e-05 | 2.60e-02 |  | 1.37 | 181 |
| GO:0008284 | Positive regulation of cell proliferation | 5.33e-05 | 2.54e-02 |  | 2.31 | 34 |
| GO:0045765 | Regulation of angiogenesis | 6.34e-05 | 2.93e-02 |  | 3.12 | 19 |
| GO:0048583 | Regulation of response to stimulus | 6.41e-05 | 2.87e-02 |  | 1.21 | 422 |
| GO:0051799 | Negative regulation of hair follicle development | 7.64e-05 | 3.32e-02 |  | 29.3 | 33 |
| GO:0006613 | Cotranslational protein targeting to membrane | 8.13e-05 | 3.44e-02 |  | 2.45 | 27 |
| GO:0051240 | Positive regulation of multicellular organismal process | 9.12e-05 | 3.75e-02 |  | 1.35 | 189 |
| GO:0044243 | Multicellular organismal catabolic process | 1.09e-04 | 4.35e-02 |  | 2.88 | 20 |
| GO:0070318 | Positive regulation of G0 to G1 transition | 1.18e-04 | 4.59e-02 |  | 12.4 | 6 |

Supplementary Table 3 Differentially expressed probes by subject age >67 years

| Transcript ID | **Gene** | **Coeff** | **p_value** | **p.adjust** |
| --- | --- | --- | --- | --- |
| ILMN_1782412 | *IRX2* | 0.3517 | 2.95E-09 | 0.0001 |
| ILMN_1691290 | *CELSR3* | 0.2296 | 2.76E-07 | 0.0058 |
| ILMN_3186390 | *LOC100128510* | -0.2183 | 1.30E-06 | 0.0182 |
| ILMN_1812824 | *SST* | -0.7052 | 1.73E-06 | 0.0183 |
| ILMN_1660292 | *MRPS21* | -0.2482 | 2.50E-06 | 0.0211 |
| ILMN_2342579 | *IL7R* | -0.4892 | 3.08E-06 | 0.0216 |
| ILMN_1691341 | *IL7R* | -0.4565 | 4.11E-06 | 0.0248 |
| ILMN_2224486 | *C3orf14* | -0.2166 | 6.92E-06 | 0.0339 |
| ILMN_1841495 | *LHFPL4* | 0.0876 | 7.23E-06 | 0.0339 |
| ILMN_1655765 | *MRPS21* | -0.1792 | 1.17E-05 | 0.0495 |

Supplementary Table 4 Differentially expressed probes with gender

| Transcript ID | Gene | Coeff | p_value | p.adjust |
| --- | --- | --- | --- | --- |
| ILMN_3238417 | *LOC100133662* | 5.25 | 3.91E-279 | 1.65E-274 |
| ILMN_1755537 | *EIF1AY* | 4.48 | 6.76E-277 | 1.43E-272 |
| ILMN_1783142 | *RPS4Y1* | 6.31 | 1.92E-275 | 2.70E-271 |
| ILMN_1670821 | *CYorf15A* | 3.12 | 2.81E-266 | 2.97E-262 |
| ILMN_2228976 | *EIF1AY* | 2.32 | 4.15E-248 | 3.50E-244 |
| ILMN_1685690 | *JARID1D* | 3.01 | 4.38E-243 | 3.08E-239 |
| ILMN_1764573 | *XIST* | -3.33 | 9.14E-232 | 5.51E-228 |
| ILMN_2191331 | *RPS4Y2* | 2.85 | 3.08E-210 | 1.62E-206 |
| ILMN_1756506 | *CYorf15B* | 1.99 | 1.30E-169 | 6.11E-166 |
| ILMN_3212921 | *LOC643123* | 1.74 | 7.30E-167 | 3.08E-163 |
| ILMN_1776195 | *TMSB4Y* | 1.8893 | 6.94E-164 | 2.66E-160 |
| ILMN_1772163 | *PRKY* | 1.7433 | 2.33E-154 | 8.18E-151 |
| ILMN_2052433 | *CYorf14* | 0.9997 | 6.56E-131 | 2.13E-127 |
| ILMN_1739587 | *UTY* | 1.0964 | 1.48E-130 | 4.47E-127 |
| ILMN_2090059 | *ZFY* | 0.9463 | 1.45E-127 | 4.07E-124 |
| ILMN_2094332 | *CYorf15B* | 1.0386 | 4.49E-118 | 1.18E-114 |
| ILMN_2143383 | *TTTY14* | 0.6197 | 2.28E-91 | 5.66E-88 |
| ILMN_1873540 | | 0.9824 | 4.15E-91 | 9.74E-88 |
| ILMN_1710136 | *HDHD1A* | -0.7164 | 4.68E-76 | 1.04E-72 |
| ILMN_2048223 | *TTTY2* | 0.4834 | 1.79E-64 | 3.78E-61 |
| ILMN_2056795 | *USP9Y* | 0.4729 | 2.98E-62 | 5.98E-59 |
| ILMN_1726816 | *LOC652814* | 0.2435 | 3.05E-50 | 5.86E-47 |
| ILMN_1716969 | *TTTY2* | 0.3111 | 3.33E-50 | 6.11E-47 |
| ILMN_1654488 | *UTX* | -0.6498 | 9.38E-50 | 1.65E-46 |
| ILMN_2166831 | *RPS4X* | -0.4743 | 3.24E-48 | 5.47E-45 |
| ILMN_2048228 | *TTTY2* | 0.3618 | 3.75E-45 | 6.08E-42 |
| ILMN_1804958 | *ZFY* | 0.2596 | 2.60E-44 | 4.07E-41 |
| ILMN_1673275 | *TRAPPC2* | -0.4179 | 4.93E-43 | 7.43E-40 |
| ILMN_2077896 | *TTTY15* | 0.2003 | 2.58E-38 | 3.75E-35 |
| ILMN_1810577 | *RPS4X* | -0.5205 | 3.85E-38 | 5.42E-35 |
| ILMN_1704431 | *LOC554203* | -0.5285 | 3.04E-34 | 4.13E-31 |
| ILMN_1684873 | *ARSD* | -0.6661 | 5.24E-33 | 6.91E-30 |
| ILMN_1687484 | *ZFX* | -0.474 | 4.73E-30 | 6.05E-27 |
| ILMN_2056032 | *CD99* | 0.3999 | 3.20E-28 | 3.97E-25 |
| ILMN_1773868 | *U2AF1L2* | -0.5428 | 1.37E-27 | 1.65E-24 |
| ILMN_2052438 | *CYorf14* | 0.2008 | 2.69E-27 | 3.15E-24 |
| ILMN_1813240 | *EIF1AX* | -0.4723 | 6.78E-26 | 7.73E-23 |
| ILMN_1664348 | *PNPLA4* | -0.3962 | 8.89E-26 | 9.87E-23 |
| ILMN_2264634 | *UTY* | 0.28 | 2.56E-25 | 2.77E-22 |
| ILMN_1684956 | *ARSD* | -0.5124 | 9.77E-25 | 1.03E-21 |
| ILMN_1786834 | *PRKX* | -0.4813 | 3.33E-24 | 3.43E-21 |
| ILMN_1727462 | *ARSE* | -0.7199 | 3.71E-24 | 3.73E-21 |
| ILMN_2210199 | *NLGN4Y* | 0.1452 | 1.10E-21 | 1.08E-18 |
| ILMN_1795243 | *LOC220433* | -0.4411 | 1.01E-20 | 9.65E-18 |
| ILMN_3280952 | *LOC391777* | -0.1513 | 1.87E-19 | 1.75E-16 |
| *ILMN_1880223* | | 0.1479 | 2.46E-18 | 2.26E-15 |
| ILMN_1794392 | *DDX3X* | -0.3169 | 2.60E-18 | 2.33E-15 |
| ILMN_3247732 | *ZRSR2* | -0.3585 | 5.45E-18 | 4.79E-15 |
| ILMN_3201216 | *LOC441550* | -0.5127 | 9.62E-18 | 8.28E-15 |
| ILMN_1755419 | *EIF1AX* | -0.3699 | 4.96E-17 | 4.18E-14 |
| ILMN_2319424 | *GYG2* | -0.4871 | 1.54E-16 | 1.27E-13 |
| ILMN_1659627 | *LOC647322* | -0.253 | 2.78E-16 | 2.25E-13 |
| ILMN_1710177 | *LOC644670* | -0.4302 | 3.07E-15 | 2.45E-12 |
| ILMN_3199929 | *LOC390183* | -0.5133 | 6.25E-15 | 4.89E-12 |
| ILMN_1665717 | *EIF2S3* | -0.1916 | 1.46E-14 | 1.12E-11 |
| ILMN_1732039 | *DDX3Y* | 0.1425 | 1.90E-14 | 1.43E-11 |
| ILMN_1703881 | *LOC441528* | -0.1421 | 2.39E-13 | 1.77E-10 |
| ILMN_3181296 | *LOC100130623* | 0.3195 | 9.34E-13 | 6.78E-10 |
| ILMN_1684017 | *GYG2* | -0.3206 | 9.48E-13 | 6.78E-10 |
| ILMN_1714523 | *HEPH* | -0.4496 | 6.47E-12 | 4.51E-09 |
| ILMN_1652006 | *SMC1A* | -0.2713 | 6.53E-12 | 4.51E-09 |
| ILMN_1778956 | *STS* | -0.1471 | 1.36E-11 | 9.24E-09 |
| ILMN_3200484 | *LOC126235* | -0.2303 | 2.10E-11 | 1.41E-08 |
| *ILMN_1882000* | | -0.2753 | 2.23E-11 | 1.47E-08 |
| ILMN_1664878 | *NLRP2* | -0.5618 | 3.48E-10 | 2.26E-07 |
| ILMN_1728540 | *FUNDC1* | -0.1905 | 4.35E-10 | 2.78E-07 |
| ILMN_3297945 | *UBA1* | -0.2446 | 4.59E-10 | 2.89E-07 |
| ILMN_1785266 | *OFD1* | -0.2817 | 1.16E-09 | 7.21E-07 |
| ILMN_3292572 | *LOC390183* | -0.2096 | 1.86E-09 | 1.13E-06 |
| ILMN_3242312 | *LOC100132864* | -0.0951 | 3.10E-09 | 1.87E-06 |
| ILMN_1798224 | *JARID1C* | -0.3163 | 6.68E-09 | 3.96E-06 |
| ILMN_3263469 | *LOC100130623* | 0.1031 | 6.76E-09 | 3.96E-06 |
| ILMN_1656165 | *USP9X* | -0.1435 | 1.13E-08 | 6.51E-06 |
| ILMN_1761456 | *ALG13* | -0.1987 | 3.33E-08 | 1.90E-05 |
| ILMN_1813893 | *DDX43* | 0.1408 | 4.50E-08 | 2.50E-05 |
| ILMN_1688318 | *MGC72104* | 0.3155 | 4.50E-08 | 2.50E-05 |
| ILMN_1760705 | *POF1B* | -0.3441 | 1.01E-07 | 5.53E-05 |
| ILMN_1662738 | *ACSM3* | 0.242 | 3.19E-07 | 0.000172 |
| ILMN_1694466 | *ZBED1* | 0.1685 | 3.66E-07 | 0.000195 |
| ILMN_1683609 | *UBE1* | -0.2665 | 5.49E-07 | 0.00029 |
| ILMN_1724341 | *CXorf45* | -0.2695 | 5.95E-07 | 0.000308 |
| ILMN_2341067 | *NLGN4X* | -0.3108 | 5.99E-07 | 0.000308 |
| ILMN_3247492 | *LOC100132373* | -0.0517 | 9.21E-07 | 0.000468 |
| ILMN_3192411 | *C10orf75* | 0.1575 | 1.14E-06 | 0.00057 |
| ILMN_1745914 | *LOC650594* | 0.0517 | 1.60E-06 | 0.00079 |
| ILMN_3249456 | *LOC728034* | -0.1414 | 1.61E-06 | 0.00079 |
| ILMN_1728011 | *NLGN4X* | -0.2004 | 2.07E-06 | 0.001005 |
| ILMN_1763326 | *C5orf25* | 0.1816 | 2.31E-06 | 0.001106 |
| ILMN_1697864 | *CXorf38* | -0.2213 | 3.03E-06 | 0.001437 |
| *ILMN_1907756* | | -0.0866 | 3.11E-06 | 0.001458 |
| ILMN_1698470 | *SYAP1* | -0.228 | 4.07E-06 | 0.001887 |
| ILMN_1776080 | *GTPBP6* | 0.1608 | 4.85E-06 | 0.002226 |
| ILMN_1672807 | *CA5B* | -0.2216 | 5.10E-06 | 0.002314 |
| ILMN_2364864 | *MB* | 0.2654 | 5.58E-06 | 0.002504 |
| ILMN_1668426 | *SPESP1* | 0.1542 | 5.73E-06 | 0.002545 |
| ILMN_3242709 | *LOC100130422* | 0.1116 | 6.90E-06 | 0.003032 |
| ILMN_3305942 | *LOC729970* | 0.197 | 8.71E-06 | 0.003788 |
| ILMN_1684922 | *LOC644322* | -0.246 | 1.28E-05 | 0.005517 |
| ILMN_3236774 | *LOC644632* | -0.1716 | 1.31E-05 | 0.005599 |
| ILMN_2308689 | *AGBL5* | 0.2293 | 1.68E-05 | 0.007079 |
| ILMN_1675939 | *IFNGR1* | -0.136 | 2.49E-05 | 0.010397 |
| ILMN_1722397 | *LOC729137* | -0.1286 | 3.73E-05 | 0.015443 |
| ILMN_3239383 | *CA5BP* | -0.0828 | 4.55E-05 | 0.018646 |
| ILMN_1755120 | *MAN1A2* | -0.1298 | 5.09E-05 | 0.020635 |
| ILMN_1774584 | *C2orf28* | 0.1074 | 5.32E-05 | 0.021374 |
| ILMN_2345119 | *NLGN4X* | -0.0768 | 5.39E-05 | 0.021439 |
| ILMN_1711810 | *PNKD* | 0.1971 | 8.02E-05 | 0.031616 |
| ILMN_3241941 | *SCARNA2* | 0.1497 | 8.66E-05 | 0.033826 |
| ILMN_3245928 | *LOC150185* | -0.0325 | 9.16E-05 | 0.035457 |
| ILMN_2140799 | *FAM24B* | 0.172 | 0.000101 | 0.038762 |
| ILMN_1801845 | *DNAL4* | 0.1218 | 0.000119 | 0.045213 |
| ILMN_1730940 | *KLHDC3* | 0.1162 | 0.00012 | 0.045213 |
| ILMN_2172755 | *DEPDC6* | 0.1809 | 0.000122 | 0.045354 |
| ILMN_2151541 | *DNAJC10* | -0.157 | 0.000123 | 0.045354 |
| ILMN_2194355 | *SOHLH2* | -0.0581 | 0.000125 | 0.045744 |
| ILMN_1701918 | *KLHDC9* | 0.1758 | 0.000134 | 0.048785 |
| ILMN_1659536 | *FAM24B* | 0.1215 | 0.000135 | 0.048785 |
| ILMN_1775170 | *MT1X* | 0.364 | 0.000137 | 0.04899 |

Supplementary Table 5 Enriched GO terms associated with male gender

| GO term | **Description** | **P-value** | **FDR q** | **Enrichment** | **Genes** |
| --- | --- | --- | --- | --- | --- |
| GO:0055114 | Oxidation-reduction process | 9.93e-08 | 1.47e-03 | 1.60 | 141 |
| GO:0006413 | Translational initiation | 3.44e-07 | 2.55e-03 | 27.7 | 6 |
| GO:0002252 | Immune effector process | 1.86e-06 | 9.20e-03 | 1.64 | 108 |
| GO:0070076 | Histone lysine demethylation | 1.94e-06 | 7.21e-03 | 75.2 | 4 |
| GO:0016577 | Histone demethylation | 2.68e-06 | 7.94e-03 | 69.9 | 4 |
| GO:0006482 | Protein demethylation | 3.60e-06 | 8.91e-03 | 65.2 | 4 |
| GO:0008214 | Protein dealkylation | 3.60e-06 | 7.64e-03 | 65.2 | 4 |
| GO:0044763 | Single-organism cellular process | 6.43e-06 | 1.19e-02 | 1.11 | 1094 |
| GO:0002263 | Cell activation involved in immune response | 8.61e-06 | 1.42e-02 | 1.84 | 67 |
| GO:0002376 | Immune system process | 1.03e-05 | 1.53e-02 | 1.37 | 210 |
| GO:0002366 | Leukocyte activation involved in immune response | 1.40e-05 | 1.88e-02 | 1.83 | 66 |
| GO:0071557 | Histone H3-K27 demethylation | 1.73e-05 | 2.14e-02 | 570. | 2 |
| GO:0071294 | Cellular response to zinc ion | 2.96e-05 | 3.38e-02 | 5.03 | 10 |
| GO:0071276 | Cellular response to cadmium ion | 3.52e-05 | 3.74e-02 | 5.10 | 11 |
| GO:0044699 | Single-organism process | 4.08e-05 | 4.04e-02 | 1.09 | 134 |
| GO:0051607 | Defense response to virus | 4.21e-05 | 3.90e-02 | 2.50 | 27 |
| GO:0070988 | Demethylation | 5.24e-05 | 4.57e-02 | 34.9 | 4 |

Differential expression was calculated using adjusted linear regression model and genes ranked by p value entered into the GO enrichment software GOrilla. FDR q-value' is the correction of the above p-value for multiple testing using the Benjamini and Hochberg method [9]. For the i^th^ term (ranked according to p-value) the FDR q-value is (p-value * number of GO terms) / i. Enrichment = (the number of genes in the intersection/ number of genes in the top of the input list)/(total number of genes associated with a specific GO term/ total number of genes).

Supplementary Table 6 Differentially expressed probes in subjects sampled under general anaesthesia

| Transcript ID | **Gene** | **coeff** | **p.adjust** |
| --- | --- | --- | --- |
| ILMN_1782412 | *IRX2* | 0.89 | 3.14E-10 |
| ILMN_2149164 | *SFRP1* | 1.28 | 2.74E-07 |
| ILMN_1723847 | *CILP* | 1.24 | 3.27E-06 |
| ILMN_1746888 | *PCOLCE2* | 1.40 | 3.27E-06 |
| ILMN_1730995 | *AFAP1L2* | 0.57 | 3.31E-06 |
| ILMN_1656369 | *C8orf4* | 1.35 | 9.51E-06 |
| ILMN_1722898 | *SFRP2* | 1.58 | 9.51E-06 |
| ILMN_3243185 | *RERGL* | 1.33 | 1.35E-05 |
| ILMN_2404917 | *AFAP1L2* | 0.58 | 1.92E-05 |
| ILMN_1770663 | *KRT24* | 1.10 | 3.59E-05 |
| ILMN_1733415 | *MFAP5* | 1.12 | 4.45E-05 |
| ILMN_2071809 | *MGP* | 1.40 | 5.96E-05 |
| ILMN_1702322 | *ALS2CL* | 0.52 | 6.06E-05 |
| ILMN_1770612 | *KRT15* | 0.74 | 6.15E-05 |
| ILMN_3241634 | *NBPF22P* | 0.35 | 6.15E-05 |
| ILMN_1752965 | *GREM1* | 1.05 | 6.73E-05 |
| ILMN_1685433 | *COL8A1* | 0.88 | 0.0001 |
| ILMN_1651958 | *MGP* | 1.35 | 0.0001 |
| ILMN_1753954 | *OLFM4* | 1.62 | 0.0001 |
| ILMN_1754103 | *CLDN11* | 0.77 | 0.0001 |
| ILMN_1796629 | *EDNRA* | 0.53 | 0.0002 |
| ILMN_1681263 | *SPINK4* | 1.77 | 0.0002 |
| ILMN_1774602 | *FBLN2* | 0.93 | 0.0002 |
| ILMN_1808114 | *LYVE1* | 1.07 | 0.0002 |
| ILMN_1713449 | *TBX3* | 0.39 | 0.0003 |
| ILMN_1676631 | *CCNO* | 0.71 | 0.0004 |
| ILMN_2167758 | *CILP* | 0.71 | 0.0004 |
| ILMN_1783846 | *RAPH1* | 0.49 | 0.0004 |
| ILMN_1745103 | *CLEC1B* | -0.23 | 0.0004 |
| ILMN_3302919 | *MYOF* | 0.57 | 0.0004 |
| ILMN_1767665 | *LOC493869* | 0.64 | 0.0004 |
| ILMN_1739496 | *PRRX1* | 0.74 | 0.0004 |
| ILMN_2102721 | *DEFA1B* | -0.46 | 0.0005 |
| ILMN_2165289 | *DEFA3* | -0.50 | 0.0005 |
| ILMN_1709486 | *SRPX* | 0.82 | 0.0006 |
| ILMN_2116877 | *OLFM4* | 1.60 | 0.0006 |
| ILMN_2390919 | *FBLN2* | 0.91 | 0.0006 |
| ILMN_1751161 | *COL7A1* | 0.64 | 0.0006 |
| ILMN_3245630 | *ECSCR* | 0.68 | 0.0006 |
| ILMN_3263575 | *C1orf81* | 0.89 | 0.0006 |
| ILMN_1762091 | *CATSPERB* | 0.43 | 0.0006 |
| ILMN_1806408 | *ACADVL* | 0.32 | 0.0006 |
| ILMN_2169383 | *REG4* | 1.63 | 0.0006 |
| ILMN_1786598 | *COL14A1* | 0.58 | 0.0006 |
| ILMN_2157240 | *MNS1* | 0.49 | 0.0006 |
| ILMN_1810289 | *FER1L3* | 0.51 | 0.0007 |
| *ILMN_1848552* | | 0.60 | 0.0007 |
| ILMN_1805448 | *EPB41L2* | 0.48 | 0.0008 |
| ILMN_1695880 | *LOX* | 0.21 | 0.0009 |
| ILMN_1790338 | *PRRX2* | 0.56 | 0.0010 |
| ILMN_3194508 | *ASAP2* | 0.48 | 0.0010 |
| ILMN_1671295 | *CCDC3* | 0.63 | 0.0010 |
| ILMN_3236259 | *PPIAL4A* | 0.34 | 0.0011 |
| ILMN_1663919 | *TFF2* | 0.82 | 0.0011 |
| ILMN_1696048 | *C13orf33* | 0.54 | 0.0011 |
| ILMN_1753347 | *DEFA4* | -0.29 | 0.0011 |
| ILMN_1791346 | *ATF3* | 0.34 | 0.0011 |
| ILMN_2389506 | *HSD11B1* | 0.28 | 0.0011 |
| ILMN_1778143 | *GRAP2* | -0.23 | 0.0012 |
| ILMN_2347145 | *DCN* | 1.02 | 0.0012 |
| ILMN_1767113 | *AOX1* | 0.55 | 0.0012 |
| ILMN_1740586 | *PLA2G2A* | 1.35 | 0.0012 |
| ILMN_1793888 | *SERPINB5* | 1.17 | 0.0012 |
| ILMN_2210753 | *XLKD1* | 0.78 | 0.0012 |
| ILMN_1790315 | *C7orf63* | 0.42 | 0.0013 |
| ILMN_2132982 | *IGFBP5* | 1.05 | 0.0014 |
| ILMN_2179717 | *C9orf61* | 0.52 | 0.0015 |
| ILMN_1792800 | *LCN6* | 0.20 | 0.0015 |
| ILMN_1678842 | *THBS2* | 0.85 | 0.0015 |
| ILMN_2317581 | *SHANK3* | 0.65 | 0.0017 |
| ILMN_1757736 | *IRX5* | 0.30 | 0.0018 |
| ILMN_3240520 | *ELTD1* | 0.40 | 0.0019 |
| ILMN_1684554 | *COL16A1* | 0.56 | 0.0020 |
| ILMN_1742534 | *COL4A5* | 0.56 | 0.0020 |
| ILMN_1755741 | *DACH1* | 0.47 | 0.0020 |
| ILMN_1707727 | *ANGPTL4* | 0.75 | 0.0022 |
| ILMN_3240524 | *MFSD6* | 0.37 | 0.0023 |
| ILMN_1681983 | *RSPO3* | 0.63 | 0.0023 |
| ILMN_1800220 | *KCTD3* | 0.42 | 0.0023 |
| ILMN_3229424 | *LOC730101* | 0.24 | 0.0024 |
| ILMN_1763941 | *LRRC49* | 0.40 | 0.0025 |
| ILMN_1653161 | *SNCG* | 0.29 | 0.0025 |
| ILMN_2315780 | *TACC2* | 0.54 | 0.0025 |
| ILMN_1745522 | *PF4V1* | -0.22 | 0.0025 |
| ILMN_1723944 | *TARP* | -0.50 | 0.0025 |
| ILMN_1798952 | *KDELR3* | 0.45 | 0.0025 |
| ILMN_3188106 | *CYTH2* | 0.50 | 0.0026 |
| ILMN_3237503 | *RNF207* | 0.37 | 0.0026 |
| ILMN_3307906 | *PALMD* | 0.47 | 0.0026 |
| ILMN_1658016 | *ZNF831* | -0.24 | 0.0027 |
| ILMN_1805543 | *ADAMTS9* | 0.49 | 0.0027 |
| ILMN_1766264 | *PI16* | 0.90 | 0.0027 |
| ILMN_2409167 | *ANXA2* | 0.29 | 0.0028 |
| ILMN_1676563 | *HTRA1* | 0.72 | 0.0028 |
| ILMN_3249167 | *SNORA63* | 0.43 | 0.0029 |
| ILMN_1706643 | *COL6A3* | 0.76 | 0.0030 |
| ILMN_1766925 | *CDH13* | 0.30 | 0.0030 |
| ILMN_1785272 | *COL1A2* | 0.84 | 0.0030 |
| ILMN_3236825 | *RAPGEF5* | 0.51 | 0.0030 |
| ILMN_1748281 | *MAPK10* | 0.40 | 0.0030 |

**Supplementary Table 7 Enriched GO terms associated with male gender**

| GO term | Description | FDR q-value | Enrichment | | Genes |
| --- | --- | --- | --- | --- | --- |
| GO:0030198 | Extracellular matrix organization | 1.27E-19 | | 5.34 | 58 |
| GO:0043062 | Extracellular structure organization | 6.34E-20 | | 5.34 | 58 |
| GO:0022610 | Biological adhesion | 1.20E-17 | | 2.05 | 198 |
| GO:0007155 | Cell adhesion | 2.62E-17 | | 2.04 | 196 |
| GO:0030155 | Regulation of cell adhesion | 1.12E-16 | | 2.71 | 113 |
| GO:0007166 | Cell surface receptor signaling pathway | 3.33E-13 | | 1.59 | 340 |
| GO:2000145 | Regulation of cell motility | 2.93E-11 | | 2.25 | 119 |
| GO:0051270 | Regulation of cellular component movement | 3.44E-11 | | 2.18 | 126 |
| GO:0006928 | Movement of cell or subcellular component | 6.19E-11 | | 1.92 | 150 |
| GO:0002376 | Immune system process | 1.60E-10 | | 1.48 | 340 |
| GO:0048583 | Regulation of response to stimulus | 1.47E-10 | | 1.34 | 537 |
| GO:0045785 | Positive regulation of cell adhesion | 1.38E-10 | | 2.70 | 78 |
| GO:0030334 | Regulation of cell migration | 1.46E-10 | | 2.26 | 111 |
| GO:0042127 | Regulation of cell proliferation | 3.26E-10 | | 1.78 | 195 |
| GO:0001775 | Cell activation | 4.81E-10 | | 1.71 | 188 |
| GO:0050776 | Regulation of immune response | 5.92E-10 | | 1.76 | 170 |
| GO:0016477 | Cell migration | 5.63E-10 | | 2.20 | 100 |
| GO:0040012 | Regulation of locomotion | 5.78E-10 | | 2.11 | 123 |
| GO:0040011 | Locomotion | 5.86E-10 | | 2.14 | 120 |
| GO:0048870 | Cell motility | 6.47E-10 | | 2.13 | 107 |
| GO:0002682 | Regulation of immune system process | 8.73E-10 | | 1.63 | 245 |
| GO:0050793 | Regulation of developmental process | 9.16E-10 | | 1.60 | 230 |
| GO:0044699 | Single-organism process | 1.67E-09 | | 1.13 | 305 |
| GO:0007165 | Signal transduction | 4.55E-09 | | 1.27 | 634 |
| GO:0044763 | Single-organism cellular process | 9.57E-09 | | 1.16 | 1025 |
| GO:0010810 | Regulation of cell-substrate adhesion | 1.22E-08 | | 3.45 | 40 |
| GO:0031589 | Cell-substrate adhesion | 2.47E-08 | | 3.53 | 37 |
| GO:0009653 | Anatomical structure morphogenesis | 2.67E-08 | | 2.11 | 94 |
| GO:0045321 | Leukocyte activation | 2.97E-08 | | 1.75 | 149 |
| GO:2000026 | Regulation of multicellular organismal development | 3.38E-08 | | 1.47 | 283 |
| GO:0098602 | Single organism cell adhesion | 4.05E-08 | | 2.30 | 70 |
| GO:0051239 | Regulation of multicellular organismal process | 4.90E-08 | | 1.36 | 402 |
| GO:0001525 | Angiogenesis | 6.23E-08 | | 3.04 | 45 |
| GO:0009611 | Response to wounding | 6.25E-08 | | 3.53 | 36 |
| GO:0044767 | Single-organism developmental process | 1.19E-07 | | 1.26 | 582 |
| GO:0032502 | Developmental process | 2.68E-07 | | 1.24 | 624 |
| GO:0022603 | Regulation of anatomical structure morphogenesis | 2.86E-07 | | 1.64 | 166 |
| GO:0045765 | Regulation of angiogenesis | 4.54E-07 | | 3.78 | 31 |
| GO:0010811 | Positive regulation of cell-substrate adhesion | 4.86E-07 | | 4.13 | 28 |
| GO:0048646 | Anatomical structure formation involved in morphogenesis | 4.92E-07 | | 3.30 | 38 |
| GO:0051241 | Negative regulation of multicellular organismal process | 1.67E-06 | | 1.67 | 145 |
| GO:0006955 | Immune response | 1.97E-06 | | 1.61 | 160 |
| GO:0048585 | Negative regulation of response to stimulus | 2.14E-06 | | 1.62 | 159 |
| GO:1901342 | Regulation of vasculature development | 2.40E-06 | | 3.21 | 35 |
| GO:0030574 | Collagen catabolic process | 3.03E-06 | | 6.39 | 17 |
| GO:0030199 | Collagen fibril organization | 2.98E-06 | | 14.9 | 10 |
| GO:0008285 | Negative regulation of cell proliferation | 3.10E-06 | | 2.42 | 56 |
| GO:0051271 | Negative regulation of cellular component movement | 3.08E-06 | | 2.39 | 53 |
| GO:0009888 | Tissue development | 4.69E-06 | | 2.09 | 73 |
| GO:0040013 | Negative regulation of locomotion | 4.76E-06 | | 3.35 | 32 |
| GO:0051093 | Negative regulation of developmental process | 5.28E-06 | | 2.21 | 65 |
| GO:0032963 | Collagen metabolic process | 5.46E-06 | | 5.76 | 18 |
| GO:0048513 | Animal organ development | 5.94E-06 | | 1.88 | 95 |
| GO:2000146 | Negative regulation of cell motility | 6.52E-06 | | 2.41 | 50 |
| GO:0007167 | Enzyme linked receptor protein signaling pathway | 6.44E-06 | | 1.69 | 123 |
| GO:0044707 | Single-multicellular organism process | 6.66E-06 | | 1.47 | 216 |
| GO:0050678 | Regulation of epithelial cell proliferation | 7.25E-06 | | 2.26 | 57 |
| GO:0044243 | Multicellular organismal catabolic process | 7.76E-06 | | 4.61 | 21 |
| GO:0009966 | Regulation of signal transduction | 1.05E-05 | | 1.38 | 283 |
| GO:0044259 | Multicellular organismal macromolecule metabolic process | 1.14E-05 | | 4.34 | 22 |
| GO:0065007 | Biological regulation | 1.14E-05 | | 1.10 | 1371 |
| GO:0051272 | Positive regulation of cellular component movement | 1.23E-05 | | 2.21 | 60 |
| GO:0022407 | Regulation of cell-cell adhesion | 1.49E-05 | | 2.14 | 63 |
| GO:0008284 | Positive regulation of cell proliferation | 1.55E-05 | | 1.68 | 122 |
| GO:0050789 | Regulation of biological process | 2.29E-05 | | 1.10 | 1306 |
| GO:1903037 | Regulation of leukocyte cell-cell adhesion | 2.40E-05 | | 2.12 | 61 |
| GO:0030336 | Negative regulation of cell migration | 3.02E-05 | | 2.17 | 54 |
| GO:0016337 | Single organismal cell-cell adhesion | 3.18E-05 | | 2.11 | 58 |
| GO:0048869 | Cellular developmental process | 3.19E-05 | | 1.61 | 135 |
| GO:0045595 | Regulation of cell differentiation | 3.19E-05 | | 1.57 | 150 |
| GO:0048584 | Positive regulation of response to stimulus | 3.29E-05 | | 1.43 | 221 |
| GO:0009887 | Animal organ morphogenesis | 4.20E-05 | | 2.68 | 39 |
| GO:0006935 | Chemotaxis | 4.39E-05 | | 2.45 | 44 |
| GO:0042330 | Taxis | 4.33E-05 | | 2.45 | 44 |
| GO:0048856 | Anatomical structure development | 4.90E-05 | | 1.49 | 175 |
| GO:0002263 | Cell activation involved in immune response | 5.11E-05 | | 1.73 | 100 |
| GO:2000147 | Positive regulation of cell motility | 5.28E-05 | | 2.15 | 57 |
| GO:2000040 | Regulation of planar cell polarity pathway involved in axis elongation | 5.45E-05 | | 2 | 2 |
| GO:2000041 | Negative regulation of planar cell polarity pathway involved in axis elongation | 5.38E-05 | | 2 | 2 |
| GO:0045055 | Regulated exocytosis | 6.76E-05 | | 1.69 | 107 |
| GO:0002684 | Positive regulation of immune system process | 6.86E-05 | | 1.68 | 112 |
| GO:0006959 | Humoral immune response | 8.41E-05 | | 3.80 | 22 |
| GO:0010646 | Regulation of cell communication | 8.51E-05 | | 1.33 | 308 |
| GO:0023051 | Regulation of signaling | 8.86E-05 | | 1.32 | 312 |
| GO:0006952 | Defense response | 8.82E-05 | | 1.49 | 170 |
| GO:0002252 | Immune effector process | 8.75E-05 | | 1.52 | 153 |
| GO:0050679 | Positive regulation of epithelial cell proliferation | 8.90E-05 | | 2.99 | 29 |
| GO:0022604 | Regulation of cell morphogenesis | 9.01E-05 | | 2.30 | 48 |
| GO:0009968 | Negative regulation of signal transduction | 9.35E-05 | | 1.61 | 127 |
| GO:0045766 | Positive regulation of angiogenesis | 9.26E-05 | | 4.31 | 20 |
| GO:0040017 | Positive regulation of locomotion | 1.02E-04 | | 2.08 | 59 |
| GO:0007169 | Transmembrane receptor protein tyrosine kinase signaling pathway | 1.10E-04 | | 1.70 | 97 |
| GO:0002274 | Myeloid leukocyte activation | 1.17E-04 | | 1.73 | 94 |
| GO:0002366 | Leukocyte activation involved in immune response | 1.17E-04 | | 1.71 | 98 |
| GO:0032101 | Regulation of response to external stimulus | 1.54E-04 | | 1.79 | 86 |
| GO:0052547 | Regulation of peptidase activity | 1.55E-04 | | 2.69 | 35 |
| GO:0051240 | Positive regulation of multicellular organismal process | 1.64E-04 | | 1.39 | 224 |
| GO:0006887 | Exocytosis | 1.76E-04 | | 1.63 | 113 |
| GO:0046903 | Secretion | 1.90E-04 | | 1.52 | 147 |
| GO:0050863 | Regulation of t cell activation | 1.98E-04 | | 2.04 | 56 |
| [GO:0048518](http://www.godatabase.org/cgi-bin/amigo/go.cgi?query=GO:0048518&view=details) | Positive regulation of biological process | 1.98E-04 | | 1.18 | 647 |
| [GO:0019730](http://www.godatabase.org/cgi-bin/amigo/go.cgi?query=GO:0019730&view=details) | Antimicrobial humoral response | 1.99E-04 | | 5.40 | 15 |
| [GO:1904018](http://www.godatabase.org/cgi-bin/amigo/go.cgi?query=GO:1904018&view=details) | Positive regulation of vasculature development | 1.98E-04 | | 3.63 | 21 |
| [GO:0034446](http://www.godatabase.org/cgi-bin/amigo/go.cgi?query=GO:0034446&view=details) | Substrate adhesion-dependent cell spreading | 2.08E-04 | | 5.17 | 14 |
| [GO:0007229](http://www.godatabase.org/cgi-bin/amigo/go.cgi?query=GO:0007229&view=details) | Integrin-mediated signaling pathway | 2.35E-04 | | 3.15 | 25 |
| [GO:0098609](http://www.godatabase.org/cgi-bin/amigo/go.cgi?query=GO:0098609&view=details) | Cell-cell adhesion | 2.69E-04 | | 1.73 | 85 |
| [GO:0044236](http://www.godatabase.org/cgi-bin/amigo/go.cgi?query=GO:0044236&view=details) | Multicellular organism metabolic process | 2.70E-04 | | 4.35 | 18 |
| [GO:0048519](http://www.godatabase.org/cgi-bin/amigo/go.cgi?query=GO:0048519&view=details) | Negative regulation of biological process | 2.69E-04 | | 1.20 | 568 |
| [GO:0002275](http://www.godatabase.org/cgi-bin/amigo/go.cgi?query=GO:0002275&view=details) | Myeloid cell activation involved in immune response | 2.83E-04 | | 1.73 | 86 |
| [GO:0022409](http://www.godatabase.org/cgi-bin/amigo/go.cgi?query=GO:0022409&view=details) | Positive regulation of cell-cell adhesion | 2.99E-04 | | 2.29 | 42 |
| [GO:0090244](http://www.godatabase.org/cgi-bin/amigo/go.cgi?query=GO:0090244&view=details) | Wnt signaling pathway involved in somitogenesis | 3.05E-04 | | 1,22 | 72 |
| [GO:0032940](http://www.godatabase.org/cgi-bin/amigo/go.cgi?query=GO:0032940&view=details) | Secretion by cell | 3.06E-04 | | 1.54 | 135 |
| [GO:0023057](http://www.godatabase.org/cgi-bin/amigo/go.cgi?query=GO:0023057&view=details) | Negative regulation of signaling | 3.21E-04 | | 1.42 | 194 |
| [GO:0032501](http://www.godatabase.org/cgi-bin/amigo/go.cgi?query=GO:0032501&view=details) | Multicellular organismal process | 3.20E-04 | | 1.46 | 160 |
| [GO:0051094](http://www.godatabase.org/cgi-bin/amigo/go.cgi?query=GO:0051094&view=details) | Positive regulation of developmental process | 3.28E-04 | | 1.59 | 119 |
| [GO:0061138](http://www.godatabase.org/cgi-bin/amigo/go.cgi?query=GO:0061138&view=details) | Morphogenesis of a branching epithelium | 3.37E-04 | | 30.4 | 5 |
| [GO:0050778](http://www.godatabase.org/cgi-bin/amigo/go.cgi?query=GO:0050778&view=details) | Positive regulation of immune response | 3.70E-04 | | 1.62 | 106 |
| [GO:0010648](http://www.godatabase.org/cgi-bin/amigo/go.cgi?query=GO:0010648&view=details) | Negative regulation of cell communication | 3.75E-04 | | 1.41 | 193 |
| [GO:0031581](http://www.godatabase.org/cgi-bin/amigo/go.cgi?query=GO:0031581&view=details) | Hemidesmosome assembly | 3.84E-04 | | 11.2 | 67 |
| [GO:0030335](http://www.godatabase.org/cgi-bin/amigo/go.cgi?query=GO:0030335&view=details) | Positive regulation of cell migration | 3.82E-04 | | 2.07 | 53 |
| [GO:0032649](http://www.godatabase.org/cgi-bin/amigo/go.cgi?query=GO:0032649&view=details) | Regulation of interferon-gamma production | 3.97E-04 | | 2.98 | 26 |
| [GO:0035295](http://www.godatabase.org/cgi-bin/amigo/go.cgi?query=GO:0035295&view=details) | Tube development | 4.02E-04 | | 2.25 | 40 |
| [GO:0044364](http://www.godatabase.org/cgi-bin/amigo/go.cgi?query=GO:0044364&view=details) | Disruption of cells of other organism | 4.39E-04 | | 7.84 | 10 |
| [GO:0031640](http://www.godatabase.org/cgi-bin/amigo/go.cgi?query=GO:0031640&view=details) | Killing of cells of other organism | 4.35E-04 | | 7.84 | 10 |
| [GO:0001763](http://www.godatabase.org/cgi-bin/amigo/go.cgi?query=GO:0001763&view=details) | Morphogenesis of a branching structure | 4.34E-04 | | 28.6 | 25 |
| [GO:0050794](http://www.godatabase.org/cgi-bin/amigo/go.cgi?query=GO:0050794&view=details) | Regulation of cellular process | 4.32E-04 | | 1.10 | 224 |
| [GO:1903039](http://www.godatabase.org/cgi-bin/amigo/go.cgi?query=GO:1903039&view=details) | Positive regulation of leukocyte cell-cell adhesion | 4.50E-04 | | 2.34 | 38 |
| [GO:0006950](http://www.godatabase.org/cgi-bin/amigo/go.cgi?query=GO:0006950&view=details) | Response to stress | 4.68E-04 | | 1.25 | 396 |
| [GO:0048523](http://www.godatabase.org/cgi-bin/amigo/go.cgi?query=GO:0048523&view=details) | Negative regulation of cellular process | 4.94E-04 | | 1.25 | 382 |
| [GO:0034330](http://www.godatabase.org/cgi-bin/amigo/go.cgi?query=GO:0034330&view=details) | Cell junction organization | 5.20E-04 | | 2.10 | 45 |
| [GO:0042119](http://www.godatabase.org/cgi-bin/amigo/go.cgi?query=GO:0042119&view=details) | Neutrophil activation | 5.35E-04 | | 1.73 | 82 |
| [GO:0007160](http://www.godatabase.org/cgi-bin/amigo/go.cgi?query=GO:0007160&view=details) | Cell-matrix adhesion | 5.53E-04 | | 3.52 | 21 |
| [GO:0042060](http://www.godatabase.org/cgi-bin/amigo/go.cgi?query=GO:0042060&view=details) | Wound healing | 5.68E-04 | | 3.97 | 18 |
| [GO:1904956](http://www.godatabase.org/cgi-bin/amigo/go.cgi?query=GO:1904956&view=details) | Regulation of midbrain dopaminergic neuron differentiation | 6.12E-04 | | 978. | 72 |
| [GO:2000051](http://www.godatabase.org/cgi-bin/amigo/go.cgi?query=GO:2000051&view=details) | Negative regulation of non-canonical wnt signaling pathway | 6.07E-04 | | 978. | 72 |
| [GO:0023056](http://www.godatabase.org/cgi-bin/amigo/go.cgi?query=GO:0023056&view=details) | Positive regulation of signaling | 6.45E-04 | | 2.24 | 43 |
| [GO:0048729](http://www.godatabase.org/cgi-bin/amigo/go.cgi?query=GO:0048729&view=details) | Tissue morphogenesis | 6.52E-04 | | 2.60 | 32 |
| [GO:0031344](http://www.godatabase.org/cgi-bin/amigo/go.cgi?query=GO:0031344&view=details) | Regulation of cell projection organization | 6.54E-04 | | 1.95 | 59 |
| [GO:1902531](http://www.godatabase.org/cgi-bin/amigo/go.cgi?query=GO:1902531&view=details) | Regulation of intracellular signal transduction | 6.51E-04 | | 1.42 | 185 |
| [GO:0080134](http://www.godatabase.org/cgi-bin/amigo/go.cgi?query=GO:0080134&view=details) | Regulation of response to stress | 6.50E-04 | | 1.52 | 135 |
| [GO:0050865](http://www.godatabase.org/cgi-bin/amigo/go.cgi?query=GO:0050865&view=details) | Regulation of cell activation | 6.46E-04 | | 1.85 | 67 |
| [GO:0032879](http://www.godatabase.org/cgi-bin/amigo/go.cgi?query=GO:0032879&view=details) | Regulation of localization | 7.20E-04 | | 1.36 | 223 |
| [GO:0002768](http://www.godatabase.org/cgi-bin/amigo/go.cgi?query=GO:0002768&view=details) | Immune response-regulating cell surface receptor signaling pathway | 7.47E-04 | | 1.82 | 65 |
| [GO:0036230](http://www.godatabase.org/cgi-bin/amigo/go.cgi?query=GO:0036230&view=details) | Granulocyte activation | 7.46E-04 | | 1.71 | 82 |
| [GO:0031347](http://www.godatabase.org/cgi-bin/amigo/go.cgi?query=GO:0031347&view=details) | Regulation of defense response | 7.50E-04 | | 1.74 | 80 |
| [GO:0043299](http://www.godatabase.org/cgi-bin/amigo/go.cgi?query=GO:0043299&view=details) | Leukocyte degranulation | 7.68E-04 | | 1.71 | 83 |
| [GO:0009967](http://www.godatabase.org/cgi-bin/amigo/go.cgi?query=GO:0009967&view=details) | Positive regulation of signal transduction | 8.45E-04 | | 1.44 | 168 |
| [GO:0060028](http://www.godatabase.org/cgi-bin/amigo/go.cgi?query=GO:0060028&view=details) | Convergent extension involved in axis elongation | 9.62E-04 | | 815 | 72 |
| [GO:0051056](http://www.godatabase.org/cgi-bin/amigo/go.cgi?query=GO:0051056&view=details) | Regulation of small gtpase mediated signal transduction | 9.61E-04 | | 2.08 | 46 |
| [GO:0043312](http://www.godatabase.org/cgi-bin/amigo/go.cgi?query=GO:0043312&view=details) | Neutrophil degranulation | 1.00E-03 | | 1.71 | 80 |
| [GO:0002283](http://www.godatabase.org/cgi-bin/amigo/go.cgi?query=GO:0002283&view=details) | Neutrophil activation involved in immune response | 1.05E-03 | | 1.71 | 80 |
| [GO:0050870](http://www.godatabase.org/cgi-bin/amigo/go.cgi?query=GO:0050870&view=details) | Positive regulation of t cell activation | 1.04E-03 | | 2.32 | 36 |
| [GO:0002683](http://www.godatabase.org/cgi-bin/amigo/go.cgi?query=GO:0002683&view=details) | Negative regulation of immune system process | 1.07E-03 | | 1.84 | 63 |
| [GO:0051336](http://www.godatabase.org/cgi-bin/amigo/go.cgi?query=GO:0051336&view=details) | Regulation of hydrolase activity | 1.15E-03 | | 1.40 | 186 |
| [GO:0050867](http://www.godatabase.org/cgi-bin/amigo/go.cgi?query=GO:0050867&view=details) | Positive regulation of cell activation | 1.15E-03 | | 2.07 | 46 |
| [GO:0046649](http://www.godatabase.org/cgi-bin/amigo/go.cgi?query=GO:0046649&view=details) | Lymphocyte activation | 1.17E-03 | | 1.84 | 61 |
| [GO:0010647](http://www.godatabase.org/cgi-bin/amigo/go.cgi?query=GO:0010647&view=details) | Positive regulation of cell communication | 1.17E-03 | | 1.42 | 170 |
| [GO:0019731](http://www.godatabase.org/cgi-bin/amigo/go.cgi?query=GO:0019731&view=details) | Antibacterial humoral response | 1.22E-03 | | 16.3 | 6 |
| [GO:0001503](http://www.godatabase.org/cgi-bin/amigo/go.cgi?query=GO:0001503&view=details) | Ossification | 1.38E-03 | | 3.81 | 18 |
| [GO:0060429](http://www.godatabase.org/cgi-bin/amigo/go.cgi?query=GO:0060429&view=details) | Epithelium development | 1.42E-03 | | 2.24 | 35 |
| [GO:0035821](http://www.godatabase.org/cgi-bin/amigo/go.cgi?query=GO:0035821&view=details) | Modification of morphology or physiology of other organism | 1.47E-03 | | 5.30 | 13 |
| [GO:0008544](http://www.godatabase.org/cgi-bin/amigo/go.cgi?query=GO:0008544&view=details) | Epidermis development | 1.50E-03 | | 4.13 | 16 |
| [GO:0016043](http://www.godatabase.org/cgi-bin/amigo/go.cgi?query=GO:0016043&view=details) | Cellular component organization | 1.52E-03 | | 1.32 | 230 |
| [GO:0002764](http://www.godatabase.org/cgi-bin/amigo/go.cgi?query=GO:0002764&view=details) | Immune response-regulating signaling pathway | 1.57E-03 | | 1.71 | 75 |
| [GO:0032102](http://www.godatabase.org/cgi-bin/amigo/go.cgi?query=GO:0032102&view=details) | Negative regulation of response to external stimulus | 1.82E-03 | | 2.69 | 27 |
| [GO:0035239](http://www.godatabase.org/cgi-bin/amigo/go.cgi?query=GO:0035239&view=details) | Tube morphogenesis | 1.83E-03 | | 20.0 | 25 |
| [GO:0031348](http://www.godatabase.org/cgi-bin/amigo/go.cgi?query=GO:0031348&view=details) | Negative regulation of defense response | 1.84E-03 | | 2.34 | 32 |
| [GO:0001894](http://www.godatabase.org/cgi-bin/amigo/go.cgi?query=GO:0001894&view=details) | Tissue homeostasis | 1.88E-03 | | 2.57 | 27 |
| [GO:1901890](http://www.godatabase.org/cgi-bin/amigo/go.cgi?query=GO:1901890&view=details) | Positive regulation of cell junction assembly | 1.89E-03 | | 14.8 | 6 |
| [GO:0050852](http://www.godatabase.org/cgi-bin/amigo/go.cgi?query=GO:0050852&view=details) | T cell receptor signaling pathway | 1.93E-03 | | 2.35 | 31 |
| [GO:0071840](http://www.godatabase.org/cgi-bin/amigo/go.cgi?query=GO:0071840&view=details) | Cellular component organization or biogenesis | 1.98E-03 | | 1.31 | 231 |
| [GO:0090288](http://www.godatabase.org/cgi-bin/amigo/go.cgi?query=GO:0090288&view=details) | Negative regulation of cellular response to growth factor stimulus | 2.05E-03 | | 9.84 | 78 |
| [GO:0045596](http://www.godatabase.org/cgi-bin/amigo/go.cgi?query=GO:0045596&view=details) | Negative regulation of cell differentiation | 2.05E-03 | | 2.07 | 46 |
| [GO:0050832](http://www.godatabase.org/cgi-bin/amigo/go.cgi?query=GO:0050832&view=details) | Defense response to fungus | 2.12E-03 | | 10.8 | 77 |
| [GO:0034329](http://www.godatabase.org/cgi-bin/amigo/go.cgi?query=GO:0034329&view=details) | Cell junction assembly | 2.13E-03 | | 2.65 | 25 |
| [GO:0050878](http://www.godatabase.org/cgi-bin/amigo/go.cgi?query=GO:0050878&view=details) | Regulation of body fluid levels | 2.13E-03 | | 1.71 | 72 |
| [GO:0002696](http://www.godatabase.org/cgi-bin/amigo/go.cgi?query=GO:0002696&view=details) | Positive regulation of leukocyte activation | 2.18E-03 | | 2.04 | 44 |
| [GO:1904338](http://www.godatabase.org/cgi-bin/amigo/go.cgi?query=GO:1904338&view=details) | Regulation of dopaminergic neuron differentiation | 2.21E-03 | | 543 | 72 |
| [GO:0051128](http://www.godatabase.org/cgi-bin/amigo/go.cgi?query=GO:0051128&view=details) | Regulation of cellular component organization | 2.22E-03 | | 1.26 | 327 |
| [GO:0032970](http://www.godatabase.org/cgi-bin/amigo/go.cgi?query=GO:0032970&view=details) | Regulation of actin filament-based process | 2.28E-03 | | 2.10 | 42 |
| [GO:0002694](http://www.godatabase.org/cgi-bin/amigo/go.cgi?query=GO:0002694&view=details) | Regulation of leukocyte activation | 2.27E-03 | | 1.63 | 84 |
| [GO:0035987](http://www.godatabase.org/cgi-bin/amigo/go.cgi?query=GO:0035987&view=details) | Endodermal cell differentiation | 2.28E-03 | | 8.17 | 58 |
| [GO:0000904](http://www.godatabase.org/cgi-bin/amigo/go.cgi?query=GO:0000904&view=details) | Cell morphogenesis involved in differentiation | 2.40E-03 | | 3.00 | 22 |
| [GO:0120035](http://www.godatabase.org/cgi-bin/amigo/go.cgi?query=GO:0120035&view=details) | Regulation of plasma membrane bounded cell projection organization | 2.51E-03 | | 1.88 | 56 |
| [GO:0070268](http://www.godatabase.org/cgi-bin/amigo/go.cgi?query=GO:0070268&view=details) | Cornification | 2.66E-03 | | 2.48 | 29 |
| [GO:0061097](http://www.godatabase.org/cgi-bin/amigo/go.cgi?query=GO:0061097&view=details) | Regulation of protein tyrosine kinase activity | 2.68E-03 | | 11.2 | 27 |
| [GO:0002253](http://www.godatabase.org/cgi-bin/amigo/go.cgi?query=GO:0002253&view=details) | Activation of immune response | 2.76E-03 | | 1.67 | 76 |
| [GO:0034332](http://www.godatabase.org/cgi-bin/amigo/go.cgi?query=GO:0034332&view=details) | Adherens junction organization | 2.83E-03 | | 2.77 | 23 |
| [GO:1901888](http://www.godatabase.org/cgi-bin/amigo/go.cgi?query=GO:1901888&view=details) | Regulation of cell junction assembly | 2.85E-03 | | 11.1 | 77 |
| [GO:0001952](http://www.godatabase.org/cgi-bin/amigo/go.cgi?query=GO:0001952&view=details) | Regulation of cell-matrix adhesion | 2.90E-03 | | 9.09 | 78 |
| [GO:0045061](http://www.godatabase.org/cgi-bin/amigo/go.cgi?query=GO:0045061&view=details) | Thymic t cell selection | 2.98E-03 | | 4.97 | 10 |
| [GO:0007044](http://www.godatabase.org/cgi-bin/amigo/go.cgi?query=GO:0007044&view=details) | Cell-substrate junction assembly | 3.09E-03 | | 6.89 | 49 |
| [GO:0030278](http://www.godatabase.org/cgi-bin/amigo/go.cgi?query=GO:0030278&view=details) | Regulation of ossification | 3.23E-03 | | 28.2 | 44 |
| [GO:0051492](http://www.godatabase.org/cgi-bin/amigo/go.cgi?query=GO:0051492&view=details) | Regulation of stress fiber assembly | 3.22E-03 | | 2.75 | 23 |
| [GO:0071481](http://www.godatabase.org/cgi-bin/amigo/go.cgi?query=GO:0071481&view=details) | Cellular response to x-ray | 3.26E-03 | | 444 | 72 |
| [GO:0060326](http://www.godatabase.org/cgi-bin/amigo/go.cgi?query=GO:0060326&view=details) | Cell chemotaxis | 3.29E-03 | | 2.50 | 27 |
| [GO:0016525](http://www.godatabase.org/cgi-bin/amigo/go.cgi?query=GO:0016525&view=details) | Negative regulation of angiogenesis | 3.35E-03 | | 5.04 | 12 |
| [GO:0021800](http://www.godatabase.org/cgi-bin/amigo/go.cgi?query=GO:0021800&view=details) | Cerebral cortex tangential migration | 3.35E-03 | | 22.2 | 44 |
| [GO:0051893](http://www.godatabase.org/cgi-bin/amigo/go.cgi?query=GO:0051893&view=details) | Regulation of focal adhesion assembly | 3.58E-03 | | 13.7 | 76 |
| [GO:0090109](http://www.godatabase.org/cgi-bin/amigo/go.cgi?query=GO:0090109&view=details) | Regulation of cell-substrate junction assembly | 3.57E-03 | | 13.7 | 76 |
| [GO:0030154](http://www.godatabase.org/cgi-bin/amigo/go.cgi?query=GO:0030154&view=details) | Cell differentiation | 3.69E-03 | | 1.31 | 242 |
| [GO:0030514](http://www.godatabase.org/cgi-bin/amigo/go.cgi?query=GO:0030514&view=details) | Negative regulation of bmp signaling pathway | 3.71E-03 | | 85.3 | 43 |
| [GO:0061844](http://www.godatabase.org/cgi-bin/amigo/go.cgi?query=GO:0061844&view=details) | Antimicrobial humoral immune response mediated by antimicrobial peptide | 3.69E-03 | | 6.71 | 9 |
| [GO:0048522](http://www.godatabase.org/cgi-bin/amigo/go.cgi?query=GO:0048522&view=details) | Positive regulation of cellular process | 3.70E-03 | | 1.16 | 613 |
| [GO:1900046](http://www.godatabase.org/cgi-bin/amigo/go.cgi?query=GO:1900046&view=details) | Regulation of hemostasis | 3.80E-03 | | 3.72 | 16 |
| [GO:0030193](http://www.godatabase.org/cgi-bin/amigo/go.cgi?query=GO:0030193&view=details) | Regulation of blood coagulation | 3.78E-03 | | 3.72 | 16 |
| [GO:2001236](http://www.godatabase.org/cgi-bin/amigo/go.cgi?query=GO:2001236&view=details) | Regulation of extrinsic apoptotic signaling pathway | 3.79E-03 | | 2.50 | 26 |
| [GO:0002429](http://www.godatabase.org/cgi-bin/amigo/go.cgi?query=GO:0002429&view=details) | Immune response-activating cell surface receptor signaling pathway | 3.99E-03 | | 1.90 | 49 |
| [GO:0071711](http://www.godatabase.org/cgi-bin/amigo/go.cgi?query=GO:0071711&view=details) | Basement membrane organization | 3.97E-03 | | 11.1 | 46 |
| [GO:0090179](http://www.godatabase.org/cgi-bin/amigo/go.cgi?query=GO:0090179&view=details) | Planar cell polarity pathway involved in neural tube closure | 4.03E-03 | | 407 | 72 |
| [GO:2000181](http://www.godatabase.org/cgi-bin/amigo/go.cgi?query=GO:2000181&view=details) | Negative regulation of blood vessel morphogenesis | 4.14E-03 | | 4.90 | 12 |
| [GO:0043508](http://www.godatabase.org/cgi-bin/amigo/go.cgi?query=GO:0043508&view=details) | Negative regulation of jun kinase activity | 4.49E-03 | | 376 | 72 |
| [GO:2000095](http://www.godatabase.org/cgi-bin/amigo/go.cgi?query=GO:2000095&view=details) | Regulation of wnt signaling pathway, planar cell polarity pathway | 4.47E-03 | | 376 | 72 |
| [GO:0060026](http://www.godatabase.org/cgi-bin/amigo/go.cgi?query=GO:0060026&view=details) | Convergent extension | 4.45E-03 | | 376 | 72 |
| [GO:0050900](http://www.godatabase.org/cgi-bin/amigo/go.cgi?query=GO:0050900&view=details) | Leukocyte migration | 4.74E-03 | | 1.82 | 53 |
| [GO:1903391](http://www.godatabase.org/cgi-bin/amigo/go.cgi?query=GO:1903391&view=details) | Regulation of adherens junction organization | 4.72E-03 | | 12.9 | 76 |
| [GO:0000902](http://www.godatabase.org/cgi-bin/amigo/go.cgi?query=GO:0000902&view=details) | Cell morphogenesis | 4.77E-03 | | 3.29 | 17 |
| [GO:0065008](http://www.godatabase.org/cgi-bin/amigo/go.cgi?query=GO:0065008&view=details) | Regulation of biological quality | 4.82E-03 | | 1.21 | 427 |
| [GO:0001101](http://www.godatabase.org/cgi-bin/amigo/go.cgi?query=GO:0001101&view=details) | Response to acid chemical | 4.85E-03 | | 1.90 | 48 |
| [GO:0007162](http://www.godatabase.org/cgi-bin/amigo/go.cgi?query=GO:0007162&view=details) | Negative regulation of cell adhesion | 4.82E-03 | | 2.04 | 39 |
| [GO:0032231](http://www.godatabase.org/cgi-bin/amigo/go.cgi?query=GO:0032231&view=details) | Regulation of actin filament bundle assembly | 4.87E-03 | | 2.56 | 25 |
| [GO:0090178](http://www.godatabase.org/cgi-bin/amigo/go.cgi?query=GO:0090178&view=details) | Regulation of establishment of planar polarity involved in neural tube closure | 5.32E-03 | | 349 | 72 |
| [GO:0043567](http://www.godatabase.org/cgi-bin/amigo/go.cgi?query=GO:0043567&view=details) | Regulation of insulin-like growth factor receptor signaling pathway | 5.58E-03 | | 27.6 | 84 |
| [GO:0045600](http://www.godatabase.org/cgi-bin/amigo/go.cgi?query=GO:0045600&view=details) | Positive regulation of fat cell differentiation | 5.63E-03 | | 30.5 | 44 |
| [GO:0051894](http://www.godatabase.org/cgi-bin/amigo/go.cgi?query=GO:0051894&view=details) | Positive regulation of focal adhesion assembly | 5.82E-03 | | 16.1 | 5 |
| [GO:0001558](http://www.godatabase.org/cgi-bin/amigo/go.cgi?query=GO:0001558&view=details) | Regulation of cell growth | 5.88E-03 | | 2.04 | 41 |
| [GO:0050730](http://www.godatabase.org/cgi-bin/amigo/go.cgi?query=GO:0050730&view=details) | Regulation of peptidyl-tyrosine phosphorylation | 6.00E-03 | | 24.3 | 44 |
| [GO:0052548](http://www.godatabase.org/cgi-bin/amigo/go.cgi?query=GO:0052548&view=details) | Regulation of endopeptidase activity | 6.15E-03 | | 2.40 | 29 |
| [GO:0050851](http://www.godatabase.org/cgi-bin/amigo/go.cgi?query=GO:0050851&view=details) | Antigen receptor-mediated signaling pathway | 6.28E-03 | | 1.98 | 39 |
| [GO:0050818](http://www.godatabase.org/cgi-bin/amigo/go.cgi?query=GO:0050818&view=details) | Regulation of coagulation | 6.46E-03 | | 3.54 | 16 |
| [GO:0002009](http://www.godatabase.org/cgi-bin/amigo/go.cgi?query=GO:0002009&view=details) | Morphogenesis of an epithelium | 6.46E-03 | | 14.9 | 25 |
| [GO:0120032](http://www.godatabase.org/cgi-bin/amigo/go.cgi?query=GO:0120032&view=details) | Regulation of plasma membrane bounded cell projection assembly | 6.57E-03 | | 2.58 | 23 |
| [GO:0033689](http://www.godatabase.org/cgi-bin/amigo/go.cgi?query=GO:0033689&view=details) | Negative regulation of osteoblast proliferation | 6.60E-03 | | 271 | 42 |
| [GO:0030168](http://www.godatabase.org/cgi-bin/amigo/go.cgi?query=GO:0030168&view=details) | Platelet activation | 6.64E-03 | | 3.61 | 16 |
| [GO:0048589](http://www.godatabase.org/cgi-bin/amigo/go.cgi?query=GO:0048589&view=details) | Developmental growth | 6.77E-03 | | 1.78 | 54 |
| [GO:0048546](http://www.godatabase.org/cgi-bin/amigo/go.cgi?query=GO:0048546&view=details) | Digestive tract morphogenesis | 7.08E-03 | | 305. | 72 |
| [GO:0009620](http://www.godatabase.org/cgi-bin/amigo/go.cgi?query=GO:0009620&view=details) | Response to fungus | 7.09E-03 | | 6.14 | 19 |
| [GO:2000080](http://www.godatabase.org/cgi-bin/amigo/go.cgi?query=GO:2000080&view=details) | Negative regulation of canonical wnt signaling pathway involved in controlling type b pancreatic cell proliferation | 7.28E-03 | | 8,56 | 21 |
| [GO:0090246](http://www.godatabase.org/cgi-bin/amigo/go.cgi?query=GO:0090246&view=details) | Convergent extension involved in somitogenesis | 7.25E-03 | | 8,56 | 21 |
| [GO:0044345](http://www.godatabase.org/cgi-bin/amigo/go.cgi?query=GO:0044345&view=details) | Stromal-epithelial cell signaling involved in prostate gland development | 7.22E-03 | | 8,56 | 21 |
| [GO:0030098](http://www.godatabase.org/cgi-bin/amigo/go.cgi?query=GO:0030098&view=details) | Lymphocyte differentiation | 7.29E-03 | | 2.17 | 32 |
| [GO:0090287](http://www.godatabase.org/cgi-bin/amigo/go.cgi?query=GO:0090287&view=details) | Regulation of cellular response to growth factor stimulus | 7.30E-03 | | 5.93 | 79 |
| [GO:0051496](http://www.godatabase.org/cgi-bin/amigo/go.cgi?query=GO:0051496&view=details) | Positive regulation of stress fiber assembly | 7.32E-03 | | 3.93 | 13 |
| [GO:0060491](http://www.godatabase.org/cgi-bin/amigo/go.cgi?query=GO:0060491&view=details) | Regulation of cell projection assembly | 7.42E-03 | | 2.54 | 23 |
| [GO:0060244](http://www.godatabase.org/cgi-bin/amigo/go.cgi?query=GO:0060244&view=details) | Negative regulation of cell proliferation involved in contact inhibition | 7.49E-03 | | 33.6 | 23 |
| [GO:0045216](http://www.godatabase.org/cgi-bin/amigo/go.cgi?query=GO:0045216&view=details) | Cell-cell junction organization | 7.48E-03 | | 1.99 | 37 |
| [GO:0048754](http://www.godatabase.org/cgi-bin/amigo/go.cgi?query=GO:0048754&view=details) | Branching morphogenesis of an epithelial tube | 7.52E-03 | | 29.1 | 24 |
| [GO:1905276](http://www.godatabase.org/cgi-bin/amigo/go.cgi?query=GO:1905276&view=details) | Regulation of epithelial tube formation | 7.58E-03 | | 287 | 72 |
| [GO:1903393](http://www.godatabase.org/cgi-bin/amigo/go.cgi?query=GO:1903393&view=details) | Positive regulation of adherens junction organization | 7.94E-03 | | 14.8 | 5 |
| [GO:0071229](http://www.godatabase.org/cgi-bin/amigo/go.cgi?query=GO:0071229&view=details) | Cellular response to acid chemical | 8.06E-03 | | 2.31 | 28 |
| [GO:0010466](http://www.godatabase.org/cgi-bin/amigo/go.cgi?query=GO:0010466&view=details) | Negative regulation of peptidase activity | 8.24E-03 | | 3.35 | 16 |
| [GO:0031175](http://www.godatabase.org/cgi-bin/amigo/go.cgi?query=GO:0031175&view=details) | Neuron projection development | 8.36E-03 | | 3.17 | 17 |
| [GO:0008360](http://www.godatabase.org/cgi-bin/amigo/go.cgi?query=GO:0008360&view=details) | Regulation of cell shape | 8.37E-03 | | 2.84 | 19 |
| [GO:0050896](http://www.godatabase.org/cgi-bin/amigo/go.cgi?query=GO:0050896&view=details) | Response to stimulus | 8.75E-03 | | 1.15 | 623 |
| [GO:0001954](http://www.godatabase.org/cgi-bin/amigo/go.cgi?query=GO:0001954&view=details) | Positive regulation of cell-matrix adhesion | 8.81E-03 | | 8.48 | 47 |
| [GO:1901343](http://www.godatabase.org/cgi-bin/amigo/go.cgi?query=GO:1901343&view=details) | Negative regulation of vasculature development | 8.79E-03 | | 4.47 | 12 |
| [GO:0050830](http://www.godatabase.org/cgi-bin/amigo/go.cgi?query=GO:0050830&view=details) | Defense response to gram-positive bacterium | 9.32E-03 | | 15.9 | 15 |
| [GO:0042110](http://www.godatabase.org/cgi-bin/amigo/go.cgi?query=GO:0042110&view=details) | T cell activation | 9.45E-03 | | 1.92 | 40 |
| [GO:0065009](http://www.godatabase.org/cgi-bin/amigo/go.cgi?query=GO:0065009&view=details) | Regulation of molecular function | 9.47E-03 | | 1.22 | 364 |
| [GO:0002727](http://www.godatabase.org/cgi-bin/amigo/go.cgi?query=GO:0002727&view=details) | Regulation of natural killer cell cytokine production | 9.78E-03 | | 107 | 92 |
| [GO:0032729](http://www.godatabase.org/cgi-bin/amigo/go.cgi?query=GO:0032729&view=details) | Positive regulation of interferon-gamma production | 9.79E-03 | | 2.98 | 18 |
| [GO:0031294](http://www.godatabase.org/cgi-bin/amigo/go.cgi?query=GO:0031294&view=details) | Lymphocyte costimulation | 9.85E-03 | | 3.35 | 16 |
| [GO:1900047](http://www.godatabase.org/cgi-bin/amigo/go.cgi?query=GO:1900047&view=details) | Negative regulation of hemostasis | 9.88E-03 | | 4.54 | 11 |
| [GO:0030195](http://www.godatabase.org/cgi-bin/amigo/go.cgi?query=GO:0030195&view=details) | Negative regulation of blood coagulation | 9.84E-03 | | 4.54 | 11 |
| [GO:0001765](http://www.godatabase.org/cgi-bin/amigo/go.cgi?query=GO:0001765&view=details) | Membrane raft assembly | 9.90E-03 | | 43.9 | 73 |
| [GO:0007528](http://www.godatabase.org/cgi-bin/amigo/go.cgi?query=GO:0007528&view=details) | Neuromuscular junction development | 9.95E-03 | | 10.0 | 26 |
| [GO:0001817](http://www.godatabase.org/cgi-bin/amigo/go.cgi?query=GO:0001817&view=details) | Regulation of cytokine production | 9.98E-03 | | 1.53 | 91 |
| [GO:2000050](http://www.godatabase.org/cgi-bin/amigo/go.cgi?query=GO:2000050&view=details) | Regulation of non-canonical wnt signaling pathway | 1.00E-02 | | 244. | 72 |
| [GO:0038065](http://www.godatabase.org/cgi-bin/amigo/go.cgi?query=GO:0038065&view=details) | Collagen-activated signaling pathway | 1.01E-02 | | 19.4 | 4 |
| [GO:0050680](http://www.godatabase.org/cgi-bin/amigo/go.cgi?query=GO:0050680&view=details) | Negative regulation of epithelial cell proliferation | 1.01E-02 | | 3.65 | 15 |
| [GO:0120036](http://www.godatabase.org/cgi-bin/amigo/go.cgi?query=GO:0120036&view=details) | Plasma membrane bounded cell projection organization | 1.03E-02 | | 1.79 | 54 |
| [GO:0060284](http://www.godatabase.org/cgi-bin/amigo/go.cgi?query=GO:0060284&view=details) | Regulation of cell development | 1.08E-02 | | 1.62 | 76 |
| [GO:0050727](http://www.godatabase.org/cgi-bin/amigo/go.cgi?query=GO:0050727&view=details) | Regulation of inflammatory response | 1.10E-02 | | 1.82 | 48 |
| [GO:0050790](http://www.godatabase.org/cgi-bin/amigo/go.cgi?query=GO:0050790&view=details) | Regulation of catalytic activity | 1.10E-02 | | 1.25 | 299 |
| [GO:0048660](http://www.godatabase.org/cgi-bin/amigo/go.cgi?query=GO:0048660&view=details) | Regulation of smooth muscle cell proliferation | 1.13E-02 | | 2.61 | 23 |
| [GO:0030036](http://www.godatabase.org/cgi-bin/amigo/go.cgi?query=GO:0030036&view=details) | Actin cytoskeleton organization | 1.16E-02 | | 1.74 | 53 |
| [GO:2000027](http://www.godatabase.org/cgi-bin/amigo/go.cgi?query=GO:2000027&view=details) | Regulation of organ morphogenesis | 1.16E-02 | | 1.82 | 46 |
| [GO:1902903](http://www.godatabase.org/cgi-bin/amigo/go.cgi?query=GO:1902903&view=details) | Regulation of supramolecular fiber organization | 1.16E-02 | | 1.72 | 55 |
| [GO:0042981](http://www.godatabase.org/cgi-bin/amigo/go.cgi?query=GO:0042981&view=details) | Regulation of apoptotic process | 1.17E-02 | | 1.33 | 191 |
| [GO:0010863](http://www.godatabase.org/cgi-bin/amigo/go.cgi?query=GO:0010863&view=details) | Positive regulation of phospholipase c activity | 1.20E-02 | | 3.78 | 12 |
| [GO:0097435](http://www.godatabase.org/cgi-bin/amigo/go.cgi?query=GO:0097435&view=details) | Supramolecular fiber organization | 1.21E-02 | | 1.95 | 40 |
| [GO:0045058](http://www.godatabase.org/cgi-bin/amigo/go.cgi?query=GO:0045058&view=details) | T cell selection | 1.22E-02 | | 4.45 | 10 |
| [GO:0072272](http://www.godatabase.org/cgi-bin/amigo/go.cgi?query=GO:0072272&view=details) | Proximal/distal pattern formation involved in metanephric nephron development | 1.21E-02 | | 8 | 11 |
| [GO:0072047](http://www.godatabase.org/cgi-bin/amigo/go.cgi?query=GO:0072047&view=details) | Proximal/distal pattern formation involved in nephron development | 1.21E-02 | | 8 | 11 |
| [GO:0001974](http://www.godatabase.org/cgi-bin/amigo/go.cgi?query=GO:0001974&view=details) | Blood vessel remodeling | 1.22E-02 | | 4.58 | 10 |
| [GO:0030029](http://www.godatabase.org/cgi-bin/amigo/go.cgi?query=GO:0030029&view=details) | Actin filament-based process | 1.25E-02 | | 1.67 | 60 |
| [GO:0032233](http://www.godatabase.org/cgi-bin/amigo/go.cgi?query=GO:0032233&view=details) | Positive regulation of actin filament bundle assembly | 1.26E-02 | | 3.50 | 14 |
| [GO:0044089](http://www.godatabase.org/cgi-bin/amigo/go.cgi?query=GO:0044089&view=details) | Positive regulation of cellular component biogenesis | 1.26E-02 | | 1.78 | 53 |
| [GO:0010941](http://www.godatabase.org/cgi-bin/amigo/go.cgi?query=GO:0010941&view=details) | Regulation of cell death | 1.27E-02 | | 1.31 | 206 |
| [GO:0016192](http://www.godatabase.org/cgi-bin/amigo/go.cgi?query=GO:0016192&view=details) | Vesicle-mediated transport | 1.28E-02 | | 1.33 | 189 |
| [GO:0010717](http://www.godatabase.org/cgi-bin/amigo/go.cgi?query=GO:0010717&view=details) | Regulation of epithelial to mesenchymal transition | 1.30E-02 | | 52.4 | 43 |
| [GO:0045597](http://www.godatabase.org/cgi-bin/amigo/go.cgi?query=GO:0045597&view=details) | Positive regulation of cell differentiation | 1.31E-02 | | 1.40 | 133 |
| [GO:0032989](http://www.godatabase.org/cgi-bin/amigo/go.cgi?query=GO:0032989&view=details) | Cellular component morphogenesis | 1.31E-02 | | 1.82 | 48 |
| [GO:0001906](http://www.godatabase.org/cgi-bin/amigo/go.cgi?query=GO:0001906&view=details) | Cell killing | 1.32E-02 | | 5.01 | 10 |
| [GO:0019065](http://www.godatabase.org/cgi-bin/amigo/go.cgi?query=GO:0019065&view=details) | Receptor-mediated endocytosis of virus by host cell | 1.35E-02 | | 97.5 | 72 |
| [GO:0075509](http://www.godatabase.org/cgi-bin/amigo/go.cgi?query=GO:0075509&view=details) | Endocytosis involved in viral entry into host cell | 1.34E-02 | | 97.5 | 72 |
| [GO:0032956](http://www.godatabase.org/cgi-bin/amigo/go.cgi?query=GO:0032956&view=details) | Regulation of actin cytoskeleton organization | 1.35E-02 | | 2.02 | 36 |
| [GO:0050817](http://www.godatabase.org/cgi-bin/amigo/go.cgi?query=GO:0050817&view=details) | Coagulation | 1.39E-02 | | 2.05 | 32 |
| [GO:0045671](http://www.godatabase.org/cgi-bin/amigo/go.cgi?query=GO:0045671&view=details) | Negative regulation of osteoclast differentiation | 1.46E-02 | | 4.91 | 59 |
| [GO:0051251](http://www.godatabase.org/cgi-bin/amigo/go.cgi?query=GO:0051251&view=details) | Positive regulation of lymphocyte activation | 1.46E-02 | | 1.94 | 38 |
| [GO:0048675](http://www.godatabase.org/cgi-bin/amigo/go.cgi?query=GO:0048675&view=details) | Axon extension | 1.48E-02 | | 7.29 | 47 |
| [GO:0048678](http://www.godatabase.org/cgi-bin/amigo/go.cgi?query=GO:0048678&view=details) | Response to axon injury | 1.50E-02 | | 6.14 | 78 |
| [GO:0051249](http://www.godatabase.org/cgi-bin/amigo/go.cgi?query=GO:0051249&view=details) | Regulation of lymphocyte activation | 1.51E-02 | | 1.59 | 71 |
| [GO:0040007](http://www.godatabase.org/cgi-bin/amigo/go.cgi?query=GO:0040007&view=details) | Growth | 1.51E-02 | | 1.65 | 61 |
| [GO:0071504](http://www.godatabase.org/cgi-bin/amigo/go.cgi?query=GO:0071504&view=details) | Cellular response to heparin | 1.51E-02 | | 9.76 | 44 |
| [GO:0002040](http://www.godatabase.org/cgi-bin/amigo/go.cgi?query=GO:0002040&view=details) | Sprouting angiogenesis | 1.52E-02 | | 4.18 | 10 |
| [GO:0010719](http://www.godatabase.org/cgi-bin/amigo/go.cgi?query=GO:0010719&view=details) | Negative regulation of epithelial to mesenchymal transition | 1.52E-02 | | 195 | 72 |
| [GO:0046851](http://www.godatabase.org/cgi-bin/amigo/go.cgi?query=GO:0046851&view=details) | Negative regulation of bone remodeling | 1.55E-02 | | 174 | 42 |
| [GO:0009628](http://www.godatabase.org/cgi-bin/amigo/go.cgi?query=GO:0009628&view=details) | Response to abiotic stimulus | 1.56E-02 | | 1.64 | 69 |
| [GO:0002757](http://www.godatabase.org/cgi-bin/amigo/go.cgi?query=GO:0002757&view=details) | Immune response-activating signal transduction | 1.59E-02 | | 1.62 | 66 |
| [GO:0050819](http://www.godatabase.org/cgi-bin/amigo/go.cgi?query=GO:0050819&view=details) | Negative regulation of coagulation | 1.60E-02 | | 4.25 | 11 |
| [GO:0002264](http://www.godatabase.org/cgi-bin/amigo/go.cgi?query=GO:0002264&view=details) | Endothelial cell activation involved in immune response | 1.67E-02 | | 2 | 61 |
| [GO:1900020](http://www.godatabase.org/cgi-bin/amigo/go.cgi?query=GO:1900020&view=details) | Positive regulation of protein kinase c activity | 1.69E-02 | | 11.2 | 24 |
| [GO:1900019](http://www.godatabase.org/cgi-bin/amigo/go.cgi?query=GO:1900019&view=details) | Regulation of protein kinase c activity | 1.69E-02 | | 11.2 | 24 |
| [GO:0014066](http://www.godatabase.org/cgi-bin/amigo/go.cgi?query=GO:0014066&view=details) | Regulation of phosphatidylinositol 3-kinase signaling | 1.69E-02 | | 2.23 | 26 |
| [GO:0030510](http://www.godatabase.org/cgi-bin/amigo/go.cgi?query=GO:0030510&view=details) | Regulation of bmp signaling pathway | 1.71E-02 | | 46.4 | 43 |
| [GO:0014745](http://www.godatabase.org/cgi-bin/amigo/go.cgi?query=GO:0014745&view=details) | Negative regulation of muscle adaptation | 1.72E-02 | | 31.4 | 43 |
| [GO:0043542](http://www.godatabase.org/cgi-bin/amigo/go.cgi?query=GO:0043542&view=details) | Endothelial cell migration | 1.74E-02 | | 4.75 | 10 |
| [GO:0021795](http://www.godatabase.org/cgi-bin/amigo/go.cgi?query=GO:0021795&view=details) | Cerebral cortex cell migration | 1.75E-02 | | 8.31 | 96 |
| [GO:0043067](http://www.godatabase.org/cgi-bin/amigo/go.cgi?query=GO:0043067&view=details) | Regulation of programmed cell death | 1.75E-02 | | 1.31 | 191 |
| [GO:0007179](http://www.godatabase.org/cgi-bin/amigo/go.cgi?query=GO:0007179&view=details) | Transforming growth factor beta receptor signaling pathway | 1.77E-02 | | 2.70 | 20 |
| [GO:1905330](http://www.godatabase.org/cgi-bin/amigo/go.cgi?query=GO:1905330&view=details) | Regulation of morphogenesis of an epithelium | 1.78E-02 | | 1.94 | 35 |
| [GO:0042476](http://www.godatabase.org/cgi-bin/amigo/go.cgi?query=GO:0042476&view=details) | Odontogenesis | 1.78E-02 | | 3.18 | 16 |
| [GO:0014068](http://www.godatabase.org/cgi-bin/amigo/go.cgi?query=GO:0014068&view=details) | Positive regulation of phosphatidylinositol 3-kinase signaling | 1.79E-02 | | 2.91 | 17 |
| [GO:0050732](http://www.godatabase.org/cgi-bin/amigo/go.cgi?query=GO:0050732&view=details) | Negative regulation of peptidyl-tyrosine phosphorylation | 1.82E-02 | | 12.5 | 25 |
| [GO:0050777](http://www.godatabase.org/cgi-bin/amigo/go.cgi?query=GO:0050777&view=details) | Negative regulation of immune response | 1.82E-02 | | 2.51 | 23 |
| [GO:0031295](http://www.godatabase.org/cgi-bin/amigo/go.cgi?query=GO:0031295&view=details) | T cell costimulation | 1.84E-02 | | 3.27 | 15 |
| [GO:0061185](http://www.godatabase.org/cgi-bin/amigo/go.cgi?query=GO:0061185&view=details) | Negative regulation of dermatome development | 1.85E-02 | | 2,44 | 71 |
| [GO:0030857](http://www.godatabase.org/cgi-bin/amigo/go.cgi?query=GO:0030857&view=details) | Negative regulation of epithelial cell differentiation | 1.85E-02 | | 4.39 | 10 |
| [GO:0031102](http://www.godatabase.org/cgi-bin/amigo/go.cgi?query=GO:0031102&view=details) | Neuron projection regeneration | 1.90E-02 | | 8.41 | 96 |
| [GO:0007178](http://www.godatabase.org/cgi-bin/amigo/go.cgi?query=GO:0007178&view=details) | Transmembrane receptor protein serine/threonine kinase signaling pathway | 1.92E-02 | | 2.15 | 28 |
| [GO:0090090](http://www.godatabase.org/cgi-bin/amigo/go.cgi?query=GO:0090090&view=details) | Negative regulation of canonical wnt signaling pathway | 1.98E-02 | | 1.98 | 33 |
| [GO:0019800](http://www.godatabase.org/cgi-bin/amigo/go.cgi?query=GO:0019800&view=details) | Peptide cross-linking via chondroitin 4-sulfate glycosaminoglycan | 1.98E-02 | | 12.1 | 84 |
| [GO:0031103](http://www.godatabase.org/cgi-bin/amigo/go.cgi?query=GO:0031103&view=details) | Axon regeneration | 2.02E-02 | | 10.7 | 45 |
| [GO:0022601](http://www.godatabase.org/cgi-bin/amigo/go.cgi?query=GO:0022601&view=details) | Menstrual cycle phase | 2.05E-02 | | 4 | 21 |
| [GO:2000054](http://www.godatabase.org/cgi-bin/amigo/go.cgi?query=GO:2000054&view=details) | Negative regulation of wnt signaling pathway involved in dorsal/ventral axis specification | 2.05E-02 | | 4 | 21 |
| [GO:2000079](http://www.godatabase.org/cgi-bin/amigo/go.cgi?query=GO:2000079&view=details) | Regulation of canonical wnt signaling pathway involved in controlling type b pancreatic cell proliferation | 2.04E-02 | | 4 | 21 |
| [GO:0014034](http://www.godatabase.org/cgi-bin/amigo/go.cgi?query=GO:0014034&view=details) | Neural crest cell fate commitment | 2.03E-02 | | 4 | 21 |
| [GO:0048865](http://www.godatabase.org/cgi-bin/amigo/go.cgi?query=GO:0048865&view=details) | Stem cell fate commitment | 2.03E-02 | | 4 | 21 |
| [GO:1900274](http://www.godatabase.org/cgi-bin/amigo/go.cgi?query=GO:1900274&view=details) | Regulation of phospholipase c activity | 2.03E-02 | | 3.55 | 12 |
| [GO:0010001](http://www.godatabase.org/cgi-bin/amigo/go.cgi?query=GO:0010001&view=details) | Glial cell differentiation | 2.04E-02 | | 3.58 | 13 |
| [GO:0010165](http://www.godatabase.org/cgi-bin/amigo/go.cgi?query=GO:0010165&view=details) | Response to x-ray | 2.06E-02 | | 163. | 72 |
| [GO:0043383](http://www.godatabase.org/cgi-bin/amigo/go.cgi?query=GO:0043383&view=details) | Negative t cell selection | 2.17E-02 | | 5.22 | 37 |
| [GO:0001932](http://www.godatabase.org/cgi-bin/amigo/go.cgi?query=GO:0001932&view=details) | Regulation of protein phosphorylation | 2.17E-02 | | 1.39 | 136 |
| [GO:0007599](http://www.godatabase.org/cgi-bin/amigo/go.cgi?query=GO:0007599&view=details) | Hemostasis | 2.23E-02 | | 2.00 | 32 |
| [GO:0001960](http://www.godatabase.org/cgi-bin/amigo/go.cgi?query=GO:0001960&view=details) | Negative regulation of cytokine-mediated signaling pathway | 2.26E-02 | | 8.68 | 26 |
| [GO:0032292](http://www.godatabase.org/cgi-bin/amigo/go.cgi?query=GO:0032292&view=details) | Peripheral nervous system axon ensheathment | 2.26E-02 | | 15.0 | 4 |
| [GO:0022011](http://www.godatabase.org/cgi-bin/amigo/go.cgi?query=GO:0022011&view=details) | Myelination in peripheral nervous system | 2.26E-02 | | 15.0 | 4 |
| [GO:0045087](http://www.godatabase.org/cgi-bin/amigo/go.cgi?query=GO:0045087&view=details) | Innate immune response | 2.27E-02 | | 1.58 | 69 |
| [GO:1902905](http://www.godatabase.org/cgi-bin/amigo/go.cgi?query=GO:1902905&view=details) | Positive regulation of supramolecular fiber organization | 2.29E-02 | | 2.18 | 26 |
| [GO:0046329](http://www.godatabase.org/cgi-bin/amigo/go.cgi?query=GO:0046329&view=details) | Negative regulation of jnk cascade | 2.29E-02 | | 152 | 72 |
| [GO:0002227](http://www.godatabase.org/cgi-bin/amigo/go.cgi?query=GO:0002227&view=details) | Innate immune response in mucosa | 2.35E-02 | | 16.3 | 4 |
| [GO:0042307](http://www.godatabase.org/cgi-bin/amigo/go.cgi?query=GO:0042307&view=details) | Positive regulation of protein import into nucleus | 2.36E-02 | | 40.3 | 43 |
| [GO:0007596](http://www.godatabase.org/cgi-bin/amigo/go.cgi?query=GO:0007596&view=details) | Blood coagulation | 2.38E-02 | | 2.01 | 31 |
| [GO:0033634](http://www.godatabase.org/cgi-bin/amigo/go.cgi?query=GO:0033634&view=details) | Positive regulation of cell-cell adhesion mediated by integrin | 2.41E-02 | | 83.9 | 22 |
| [GO:0050768](http://www.godatabase.org/cgi-bin/amigo/go.cgi?query=GO:0050768&view=details) | Negative regulation of neurogenesis | 2.46E-02 | | 2.10 | 29 |
| [GO:0010951](http://www.godatabase.org/cgi-bin/amigo/go.cgi?query=GO:0010951&view=details) | Negative regulation of endopeptidase activity | 2.51E-02 | | 2.35 | 23 |
| [GO:0060828](http://www.godatabase.org/cgi-bin/amigo/go.cgi?query=GO:0060828&view=details) | Regulation of canonical wnt signaling pathway | 2.50E-02 | | 1.78 | 43 |
| [GO:0030030](http://www.godatabase.org/cgi-bin/amigo/go.cgi?query=GO:0030030&view=details) | Cell projection organization | 2.53E-02 | | 1.61 | 67 |
| [GO:0006968](http://www.godatabase.org/cgi-bin/amigo/go.cgi?query=GO:0006968&view=details) | Cellular defense response | 2.53E-02 | | 2.66 | 18 |
| [GO:1904591](http://www.godatabase.org/cgi-bin/amigo/go.cgi?query=GO:1904591&view=details) | Positive regulation of protein import | 2.52E-02 | | 39.4 | 43 |
| [GO:0031953](http://www.godatabase.org/cgi-bin/amigo/go.cgi?query=GO:0031953&view=details) | Negative regulation of protein autophosphorylation | 2.57E-02 | | 27.4 | 73 |
| [GO:0034104](http://www.godatabase.org/cgi-bin/amigo/go.cgi?query=GO:0034104&view=details) | Negative regulation of tissue remodeling | 2.62E-02 | | 128 | 42 |
| [GO:0090101](http://www.godatabase.org/cgi-bin/amigo/go.cgi?query=GO:0090101&view=details) | Negative regulation of transmembrane receptor protein serine/threonine kinase signaling pathway | 2.62E-02 | | 8.96 | 76 |
| [GO:0022029](http://www.godatabase.org/cgi-bin/amigo/go.cgi?query=GO:0022029&view=details) | Telencephalon cell migration | 2.62E-02 | | 6.36 | 97 |
| [GO:0048010](http://www.godatabase.org/cgi-bin/amigo/go.cgi?query=GO:0048010&view=details) | Vascular endothelial growth factor receptor signaling pathway | 2.64E-02 | | 2.73 | 18 |
| [GO:0048662](http://www.godatabase.org/cgi-bin/amigo/go.cgi?query=GO:0048662&view=details) | Negative regulation of smooth muscle cell proliferation | 2.63E-02 | | 3.78 | 11 |
| [GO:0050673](http://www.godatabase.org/cgi-bin/amigo/go.cgi?query=GO:0050673&view=details) | Epithelial cell proliferation | 2.66E-02 | | 2.87 | 17 |
| [GO:0030324](http://www.godatabase.org/cgi-bin/amigo/go.cgi?query=GO:0030324&view=details) | Lung development | 2.69E-02 | | 3.31 | 14 |
| [GO:0061299](http://www.godatabase.org/cgi-bin/amigo/go.cgi?query=GO:0061299&view=details) | Retina vasculature morphogenesis in camera-type eye | 2.68E-02 | | 12.8 | 44 |
| [GO:0010721](http://www.godatabase.org/cgi-bin/amigo/go.cgi?query=GO:0010721&view=details) | Negative regulation of cell development | 2.70E-02 | | 2.00 | 32 |
| [GO:0043569](http://www.godatabase.org/cgi-bin/amigo/go.cgi?query=GO:0043569&view=details) | Negative regulation of insulin-like growth factor receptor signaling pathway | 2.77E-02 | | 92.3 | 32 |
| [GO:0072376](http://www.godatabase.org/cgi-bin/amigo/go.cgi?query=GO:0072376&view=details) | Protein activation cascade | 2.77E-02 | | 4.45 | 10 |
| [GO:0022617](http://www.godatabase.org/cgi-bin/amigo/go.cgi?query=GO:0022617&view=details) | Extracellular matrix disassembly | 2.79E-02 | | 4.10 | 11 |
| [GO:0043409](http://www.godatabase.org/cgi-bin/amigo/go.cgi?query=GO:0043409&view=details) | Negative regulation of mapk cascade | 2.85E-02 | | 5.05 | 78 |
| [GO:0033630](http://www.godatabase.org/cgi-bin/amigo/go.cgi?query=GO:0033630&view=details) | Positive regulation of cell adhesion mediated by integrin | 2.87E-02 | | 29.6 | 23 |
| [GO:0050920](http://www.godatabase.org/cgi-bin/amigo/go.cgi?query=GO:0050920&view=details) | Regulation of chemotaxis | 2.87E-02 | | 2.04 | 29 |
| [GO:0010562](http://www.godatabase.org/cgi-bin/amigo/go.cgi?query=GO:0010562&view=details) | Positive regulation of phosphorus metabolic process | 2.90E-02 | | 1.45 | 102 |
| [GO:0045937](http://www.godatabase.org/cgi-bin/amigo/go.cgi?query=GO:0045937&view=details) | Positive regulation of phosphate metabolic process | 2.90E-02 | | 1.45 | 102 |
| [GO:0032682](http://www.godatabase.org/cgi-bin/amigo/go.cgi?query=GO:0032682&view=details) | Negative regulation of chemokine production | 2.97E-02 | | 29.0 | 43 |
| [GO:0060055](http://www.godatabase.org/cgi-bin/amigo/go.cgi?query=GO:0060055&view=details) | Angiogenesis involved in wound healing | 2.97E-02 | | 6.50 | 6 |
| [GO:0051259](http://www.godatabase.org/cgi-bin/amigo/go.cgi?query=GO:0051259&view=details) | Protein oligomerization | 2.98E-02 | | 2.83 | 17 |
| [GO:0043408](http://www.godatabase.org/cgi-bin/amigo/go.cgi?query=GO:0043408&view=details) | Regulation of mapk cascade | 2.99E-02 | | 1.58 | 70 |
| [GO:0001656](http://www.godatabase.org/cgi-bin/amigo/go.cgi?query=GO:0001656&view=details) | Metanephros development | 3.01E-02 | | 4.39 | 59 |
| [GO:0051130](http://www.godatabase.org/cgi-bin/amigo/go.cgi?query=GO:0051130&view=details) | Positive regulation of cellular component organization | 3.05E-02 | | 1.31 | 174 |
| [GO:0035567](http://www.godatabase.org/cgi-bin/amigo/go.cgi?query=GO:0035567&view=details) | Non-canonical wnt signaling pathway | 3.04E-02 | | 2.02 | 28 |
| [GO:0045859](http://www.godatabase.org/cgi-bin/amigo/go.cgi?query=GO:0045859&view=details) | Regulation of protein kinase activity | 3.06E-02 | | 13.7 | 74 |
| [GO:0030162](http://www.godatabase.org/cgi-bin/amigo/go.cgi?query=GO:0030162&view=details) | Regulation of proteolysis | 3.08E-02 | | 2.35 | 24 |
| [GO:0051897](http://www.godatabase.org/cgi-bin/amigo/go.cgi?query=GO:0051897&view=details) | Positive regulation of protein kinase b signaling | 3.12E-02 | | 2.82 | 17 |
| [GO:0043085](http://www.godatabase.org/cgi-bin/amigo/go.cgi?query=GO:0043085&view=details) | Positive regulation of catalytic activity | 3.13E-02 | | 5.02 | 88 |
| [GO:0061045](http://www.godatabase.org/cgi-bin/amigo/go.cgi?query=GO:0061045&view=details) | Negative regulation of wound healing | 3.14E-02 | | 3.63 | 12 |
| [GO:0033688](http://www.godatabase.org/cgi-bin/amigo/go.cgi?query=GO:0033688&view=details) | Regulation of osteoblast proliferation | 3.17E-02 | | 14.9 | 94 |
| [GO:0060761](http://www.godatabase.org/cgi-bin/amigo/go.cgi?query=GO:0060761&view=details) | Negative regulation of response to cytokine stimulus | 3.17E-02 | | 7.92 | 26 |
| [GO:0051345](http://www.godatabase.org/cgi-bin/amigo/go.cgi?query=GO:0051345&view=details) | Positive regulation of hydrolase activity | 3.16E-02 | | 1.38 | 129 |
| [GO:0072268](http://www.godatabase.org/cgi-bin/amigo/go.cgi?query=GO:0072268&view=details) | Pattern specification involved in metanephros development | 3.20E-02 | | 4 | 11 |
| [GO:0072086](http://www.godatabase.org/cgi-bin/amigo/go.cgi?query=GO:0072086&view=details) | Specification of loop of henle identity | 3.19E-02 | | 4 | 11 |
| [GO:0051248](http://www.godatabase.org/cgi-bin/amigo/go.cgi?query=GO:0051248&view=details) | Negative regulation of protein metabolic process | 3.19E-02 | | 2.03 | 33 |
| [GO:1902533](http://www.godatabase.org/cgi-bin/amigo/go.cgi?query=GO:1902533&view=details) | Positive regulation of intracellular signal transduction | 3.24E-02 | | 1.45 | 99 |
| [GO:0044087](http://www.godatabase.org/cgi-bin/amigo/go.cgi?query=GO:0044087&view=details) | Regulation of cellular component biogenesis | 3.33E-02 | | 1.40 | 116 |
| [GO:0070836](http://www.godatabase.org/cgi-bin/amigo/go.cgi?query=GO:0070836&view=details) | Caveola assembly | 3.43E-02 | | 73.1 | 72 |
| [GO:0072009](http://www.godatabase.org/cgi-bin/amigo/go.cgi?query=GO:0072009&view=details) | Nephron epithelium development | 3.42E-02 | | 9.27 | 25 |
| [GO:0097102](http://www.godatabase.org/cgi-bin/amigo/go.cgi?query=GO:0097102&view=details) | Endothelial tip cell fate specification | 3.42E-02 | | 45.9 | 32 |
| [GO:0010942](http://www.godatabase.org/cgi-bin/amigo/go.cgi?query=GO:0010942&view=details) | Positive regulation of cell death | 3.45E-02 | | 1.45 | 94 |
| [GO:0031069](http://www.godatabase.org/cgi-bin/amigo/go.cgi?query=GO:0031069&view=details) | Hair follicle morphogenesis | 3.45E-02 | | 5.57 | 87 |
| [GO:0032873](http://www.godatabase.org/cgi-bin/amigo/go.cgi?query=GO:0032873&view=details) | Negative regulation of stress-activated mapk cascade | 3.47E-02 | | 119 | 72 |
| [GO:0070303](http://www.godatabase.org/cgi-bin/amigo/go.cgi?query=GO:0070303&view=details) | Negative regulation of stress-activated protein kinase signaling cascade | 3.46E-02 | | 119 | 72 |
| [GO:0032331](http://www.godatabase.org/cgi-bin/amigo/go.cgi?query=GO:0032331&view=details) | Negative regulation of chondrocyte differentiation | 3.52E-02 | | 4.51 | 8 |
| [GO:0042221](http://www.godatabase.org/cgi-bin/amigo/go.cgi?query=GO:0042221&view=details) | Response to chemical | 3.55E-02 | | 1.21 | 315 |
| [GO:0045665](http://www.godatabase.org/cgi-bin/amigo/go.cgi?query=GO:0045665&view=details) | Negative regulation of neuron differentiation | 3.56E-02 | | 2.19 | 24 |
| [GO:0061041](http://www.godatabase.org/cgi-bin/amigo/go.cgi?query=GO:0061041&view=details) | Regulation of wound healing | 3.56E-02 | | 3.00 | 16 |

**Supplementary Table 8 Differentially expressed genes by colorectal side with fold-change >2**

| Transcript ID | Gene | FC | p_value | p.adjust |
| --- | --- | --- | --- | --- |
| ILMN_1801832 | *PRAC* | 0.033 | 1.77E-83 | 4.46E-79 |
| ILMN_3248384 | *PRAC* | 0.034 | 2.11E-83 | 4.46E-79 |
| ILMN_2391400 | *PITX2* | 4.734 | 3.24E-63 | 4.56E-59 |
| ILMN_1796847 | *PITX2* | 2.326 | 2.67E-51 | 2.81E-47 |
| ILMN_1742677 | *HOXB13* | 0.220 | 4.00E-48 | 3.37E-44 |
| ILMN_2204545 | *ST3GAL4* | 0.368 | 1.25E-33 | 8.79E-30 |
| ILMN_1691736 | *ST6GALNAC6* | 0.416 | 3.07E-33 | 1.77E-29 |
| ILMN_1769839 | *L1TD1* | 2.585 | 3.36E-33 | 1.77E-29 |
| ILMN_1746676 | *CLDN8* | 0.258 | 6.05E-31 | 2.84E-27 |
| ILMN_2072568 | *CLDN8* | 0.238 | 8.37E-31 | 3.21E-27 |
| ILMN_1791890 | *SPON1* | 0.455 | 1.63E-29 | 5.28E-26 |
| ILMN_1666109 | *MB* | 2.119 | 4.34E-29 | 1.31E-25 |
| ILMN_1696028 | *ETNK1* | 2.207 | 4.20E-28 | 1.18E-24 |
| ILMN_1758039 | *KRTAP13-2* | 0.329 | 6.58E-25 | 1.46E-21 |
| ILMN_2316778 | *ETNK1* | 2.019 | 8.00E-25 | 1.69E-21 |
| ILMN_1734694 | *MEP1B* | 2.077 | 1.17E-24 | 2.36E-21 |
| ILMN_1687306 | *LGALS2* | 0.464 | 3.60E-22 | 5.63E-19 |
| ILMN_1678478 | *CHST5* | 0.497 | 2.45E-21 | 3.69E-18 |
| ILMN_1677108 | *CAPN13* | 0.440 | 4.01E-21 | 5.32E-18 |
| ILMN_1671478 | *CKB* | 0.405 | 5.31E-21 | 6.78E-18 |
| ILMN_2355486 | *FAM3B* | 2.229 | 9.61E-21 | 1.19E-17 |
| ILMN_3239925 | *LOC25845* | 2.506 | 2.00E-20 | 2.22E-17 |
| ILMN_1775814 | *GHR* | 0.492 | 6.56E-20 | 6.92E-17 |
| *ILMN_1836218* | | 2.510 | 3.62E-19 | 3.73E-16 |
| ILMN_1725387 | *TMEM200A* | 0.484 | 9.77E-19 | 8.96E-16 |
| ILMN_3307693 | *WFDC2* | 0.406 | 3.12E-18 | 2.80E-15 |
| ILMN_1751227 | *LOC401321* | 0.495 | 5.43E-18 | 4.58E-15 |
| ILMN_2382679 | *REG3A* | 2.491 | 6.45E-18 | 5.13E-15 |
| ILMN_1704376 | *GLDN* | 0.384 | 9.02E-17 | 5.94E-14 |
| ILMN_1753954 | *OLFM4* | 3.004 | 1.82E-16 | 1.15E-13 |
| ILMN_1770424 | *DEFA5* | 3.775 | 2.48E-16 | 1.54E-13 |
| ILMN_2116877 | *OLFM4* | 3.073 | 2.22E-15 | 1.28E-12 |
| ILMN_1724375 | *MUC17* | 0.443 | 2.38E-15 | 1.34E-12 |
| ILMN_1690017 | *SPINK5* | 0.462 | 4.40E-14 | 2.00E-11 |
| ILMN_1722489 | *TFF1* | 0.401 | 8.66E-13 | 2.90E-10 |
| ILMN_1759089 | *DEFA6* | 2.401 | 7.27E-12 | 2.05E-09 |
| ILMN_1763837 | *ANPEP* | 2.144 | 1.86E-11 | 4.50E-09 |
| ILMN_1693192 | *PI3* | 0.444 | 3.04E-11 | 7.09E-09 |
| ILMN_2060578 | *INSL5* | 0.433 | 4.89E-11 | 1.10E-08 |
| ILMN_1802441 | *REG1A* | 2.411 | 9.87E-11 | 2.06E-08 |
| ILMN_1703075 | *PYY* | 0.489 | 1.45E-10 | 2.89E-08 |
| ILMN_1799020 | *MUC12* | 0.465 | 1.89E-10 | 3.63E-08 |
| ILMN_1724396 | *GCG* | 0.489 | 1.95E-10 | 3.72E-08 |
| ILMN_1679357 | *DEFA1B* | 2.069 | 3.55E-10 | 6.34E-08 |
| ILMN_1789096 | *OSTalpha* | 2.098 | 2.63E-09 | 3.74E-07 |
| ILMN_1801886 | *SLC28A2* | 0.476 | 2.03E-07 | 1.75E-05 |
| ILMN_1751607 | *FOSB* | 2.155 | 8.21E-07 | 5.60E-05 |

**Supplementary Table 9 Genes reported to be differentially expressed in stroma and epithelium compared between stripped mucosa and biopsy samples in current dataset**

| Probe ID | Gene | Logfc | Avr. Exp | p-value | Compartment |
| --- | --- | --- | --- | --- | --- |
| ILMN_1704730 | *CD93* | 0.20 | 8.47 | 0.01 | Stromal |
| ILMN_1688480 | *CCND1* | -0.22 | 10.13 | 0.01 | Epithelial |
| ILMN_2229877 | *PCDH18* | -0.17 | 8.29 | 0.05 | Stromal |
| ILMN_2058251 | *VIM* | 0.12 | 11.02 | 0.07 | Stromal |
| ILMN_3249032 | *EPCAM* | -0.15 | 13.20 | 0.08 | Epithelial |
| ILMN_1796801 | *ABCA8* | -0.13 | 8.63 | 0.11 | Stromal |
| ILMN_1727087 | *GJA1* | -0.12 | 7.23 | 0.12 | Stromal |
| ILMN_1671703 | *ACTA2* | -0.14 | 12.24 | 0.14 | Stromal |
| ILMN_1782538 | *VIM* | 0.10 | 11.83 | 0.14 | Stromal |
| ILMN_2196328 | *POSTN* | -0.09 | 6.56 | 0.21 | Stromal |

Unadjusted P value given. As expression of 42 genes was compared, level of statistical significance was set at 0.001

**Supplementary Figure 5 Candidate gene expression in matched biopsies and stripped mucosa samples**

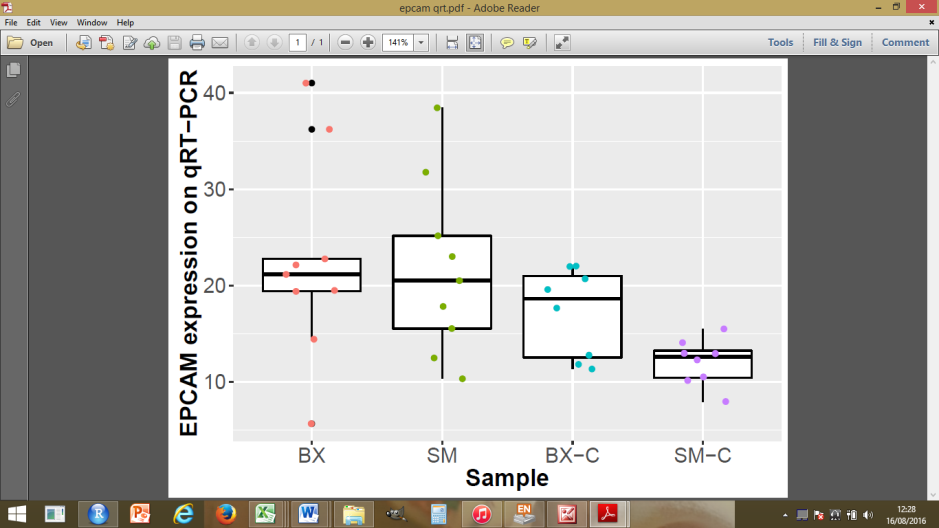

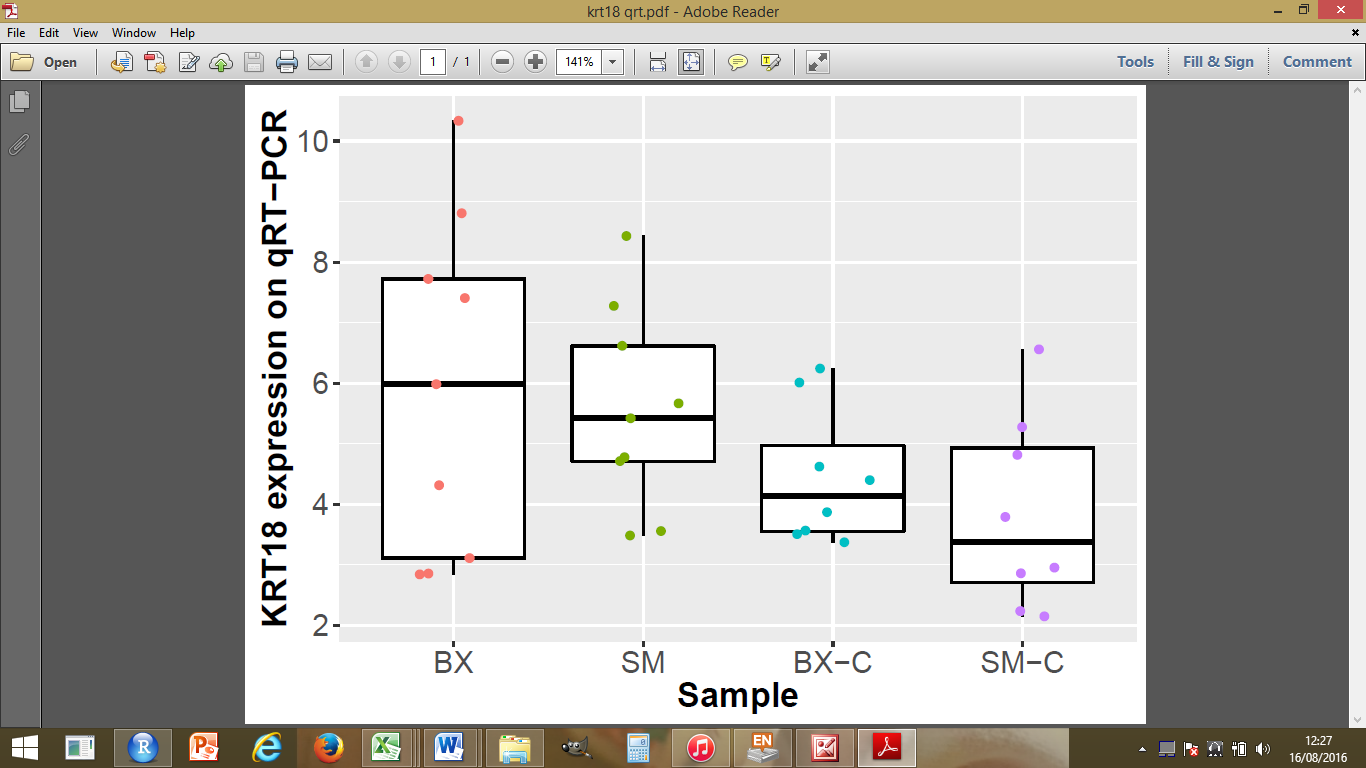

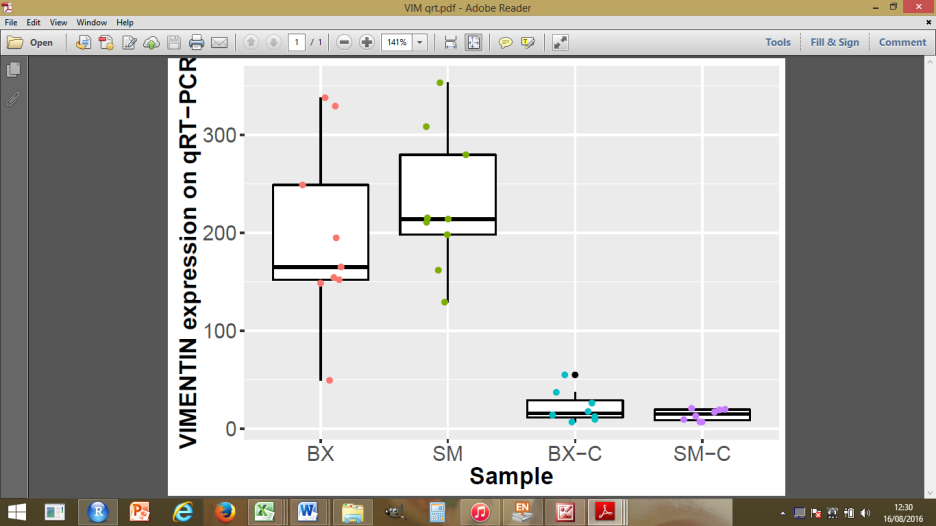

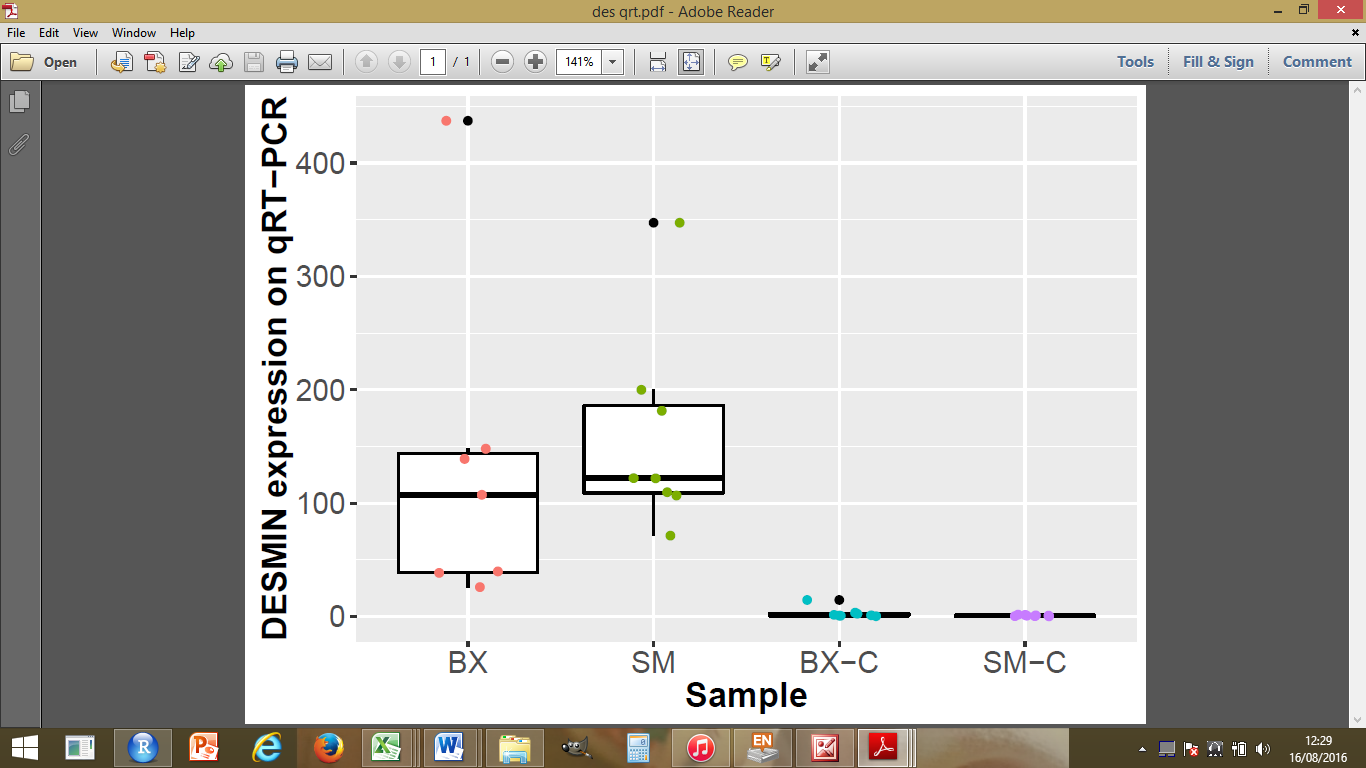

Matched biopsy and stripped mucosa samples were taken in 9 patients. The matched samples together with biological replicates were included for analysis on qRT-PCR. BX – biopsy, SM- stripped mucosa, No significant differences in candidate gene expression was seen (p>0.05).

**Supplementary Table 10 Enriched GO terms associated with delay to RNA preservation**

| GO term | Description | FDR q-value | Enrichment | Genes |
| --- | --- | --- | --- | --- |
| GO:0031325 | Positive regulation of cellular metabolic process | 1.95e-03 | 1.30 | 384 |
| GO:0009893 | Positive regulation of metabolic process | 1.03e-03 | 1.28 | 410 |
| GO:0010604 | Positive regulation of macromolecule metabolic process | 1.85e-03 | 1.29 | 382 |
| GO:0051173 | Positive regulation of nitrogen compound metabolic process | 2.38e-03 | 1.29 | 358 |
| GO:0044238 | Primary metabolic process | 6.10e-03 | 1.16 | 653 |
| GO:0008152 | Metabolic process | 5.93e-03 | 1.12 | 934 |
| GO:0071704 | Organic substance metabolic process | 5.09e-03 | 1.15 | 679 |
| GO:0009611 | Response to wounding | 5.00e-03 | 14.30 | 7 |
| GO:0006915 | Apoptotic process | 7.98e-03 | 2.13 | 9 |
| GO:0048513 | Animal organ development | 1.23e-02 | 8.82 | 8 |
| GO:0043122 | Regulation of I-kappab kinase/NF-kappab signaling | 1.32e-02 | 2.00 | 9 |
| GO:0043170 | Macromolecule metabolic process | 2.14e-02 | 1.17 | 563 |
| GO:0031960 | Response to corticosteroid | 2.16e-02 | 5.64 | 1 |
| GO:0006807 | Nitrogen compound metabolic process | 2.23e-02 | 1.15 | 611 |
| GO:0019222 | Regulation of metabolic process | 2.48e-02 | 1.14 | 801 |
| GO:0009891 | Positive regulation of biosynthetic process | 2.35e-02 | 1.35 | 217 |
| GO:0010557 | Positive regulation of macromolecule biosynthetic process | 2.22e-02 | 1.36 | 201 |
| GO:0060527 | Prostate epithelial cord arborization involved in prostate glandular acinus morphogenesis | 2.57e-02 | 439.00 | 2 |
| GO:0048519 | Negative regulation of biological process | 2.82e-02 | 1.20 | 476 |
| GO:0031328 | Positive regulation of cellular biosynthetic process | 2.69e-02 | 1.34 | 213 |
| GO:0031099 | Regeneration | 2.81e-02 | 38.80 | 4 |
| GO:0010628 | Positive regulation of gene expression | 2.79e-02 | 1.33 | 226 |
| GO:0051384 | Response to glucocorticoid | 2.68e-02 | 5.72 | 0 |
| GO:0031326 | Regulation of cellular biosynthetic process | 2.62e-02 | 1.18 | 532 |
| GO:0044237 | Cellular metabolic process | 2.82e-02 | 1.12 | 853 |
| GO:0031323 | Regulation of cellular metabolic process | 2.79e-02 | 1.15 | 698 |
| GO:0010468 | Regulation of gene expression | 2.82e-02 | 1.17 | 572 |
| GO:0009889 | Regulation of biosynthetic process | 2.90e-02 | 1.18 | 539 |
| GO:0044710 | Single-organism metabolic process | 3.15e-02 | 1.20 | 468 |
| GO:0051254 | Positive regulation of RNA metabolic process | 3.06e-02 | 1.36 | 189 |
| GO:0065009 | Regulation of molecular function | 3.18e-02 | 1.43 | 140 |
| GO:0010556 | Regulation of macromolecule biosynthetic process | 3.21e-02 | 1.18 | 510 |
| GO:0033993 | Response to lipid | 3.36e-02 | 10.80 | 6 |
| GO:0070482 | Response to oxygen levels | 3.44e-02 | 1.99 | 2 |
| GO:0051247 | Positive regulation of protein metabolic process | 3.39e-02 | 1.33 | 210 |
| GO:0014070 | Response to organic cyclic compound | 3.47e-02 | 10.60 | 6 |
| GO:0010941 | Regulation of cell death | 3.43e-02 | 1.35 | 188 |
| GO:0060442 | Branching involved in prostate gland morphogenesis | 3.51e-02 | 292.00 | 2 |
| GO:0048523 | Negative regulation of cellular process | 3.53e-02 | 1.20 | 430 |
| GO:0080090 | Regulation of primary metabolic process | 3.47e-02 | 1.14 | 683 |
| GO:0044092 | Negative regulation of molecular function | 3.51e-02 | 2.00 | 4 |
| GO:0048545 | Response to steroid hormone | 3.47e-02 | 4.41 | 2 |
| GO:0045765 | Regulation of angiogenesis | 3.47e-02 | 2.06 | 7 |
| GO:0060255 | Regulation of macromolecule metabolic process | 3.60e-02 | 1.14 | 687 |
| GO:0050793 | Regulation of developmental process | 3.56e-02 | 1.28 | 265 |
| GO:0012501 | Programmed cell death | 3.53e-02 | 1.86 | 2 |
| GO:0060734 | Regulation of endoplasmic reticulum stress-induced eif2 alpha phosphorylation | 3.50e-02 | 133.00 | 2 |
| GO:0021520 | Spinal cord motor neuron cell fate specification | 3.45e-02 | 244.00 | 2 |
| GO:0031100 | Animal organ regeneration | 3.57e-02 | 28.80 | 4 |
| GO:0042981 | Regulation of apoptotic process | 3.56e-02 | 1.36 | 175 |
| GO:0050728 | Negative regulation of inflammatory response | 3.94e-02 | 3.98 | 14 |
| GO:0045935 | Positive regulation of nucleobase-containing compound metabolic process | 4.01e-02 | 1.31 | 215 |
| GO:0043542 | Endothelial cell migration | 3.95e-02 | 5.37 | 110 |
| GO:0043123 | Positive regulation of I-kappab kinase/NF-kappab signaling | 3.94e-02 | 1.99 | 737 |
| GO:0032270 | Positive regulation of cellular protein metabolic process | 3.98e-02 | 1.33 | 196 |
| GO:0010888 | Negative regulation of lipid storage | 4.12e-02 | 5.74 | 48 |
| GO:0071403 | Cellular response to high density lipoprotein particle stimulus | 4.56e-02 | 5709.00 | 31 |
| GO:0033552 | Response to vitamin B3 | 4.48e-02 | 5709.00 | 31 |
| GO:0048518 | Positive regulation of biological process | 4.50e-02 | 1.16 | 576 |
| GO:0042221 | Response to chemical | 4.66e-02 | 1.44 | 122 |
| GO:1901342 | Regulation of vasculature development | 4.83e-02 | 1.95 | 339 |
| GO:1904659 | Glucose transmembrane transport | 4.90e-02 | 47.50 | 603 |
| GO:0035428 | Hexose transmembrane transport | 4.82e-02 | 47.50 | 603 |
| GO:1905950 | Monosaccharide transmembrane transport | 4.74e-02 | 47.50 | 603 |
| GO:0008219 | Cell death | 4.71e-02 | 1.81 | 853 |
| GO:0030522 | Intracellular receptor signaling pathway | 4.71e-02 | 2.00 | 635 |
| GO:0036293 | Response to decreased oxygen levels | 4.80e-02 | 1.96 | 139 |
| GO:0043067 | Regulation of programmed cell death | 4.74e-02 | 1.35 | 175 |
| GO:0009719 | Response to endogenous stimulus | 4.99e-02 | 1.81 | 752 |
| GO:0071347 | Cellular response to interleukin-1 | 4.99e-02 | 53.90 | 143 |
| GO:0031348 | Negative regulation of defense response | 4.96e-02 | 3.11 | 16 |
| GO:0002548 | Monocyte chemotaxis | 4.96e-02 | 225.00 | 2 |
